# Applicability of subcortical EEG metrics of synaptopathy to older listeners with impaired audiograms

**DOI:** 10.1101/479246

**Authors:** Markus Garrett, Sarah Verhulst

**Affiliations:** Medizinische Physik and Cluster of Excellence “Hearing4all”, Department of Medical Physics and Acoustics, University of Oldenburg, Oldenburg, Germany; Hearing Technology @ WAVES, Department of Information Technology, Ghent University, Ghent, Belgium

**Keywords:** auditory brainstem response, envelope following response, cochlear synaptopathy, diagnostics, sensorineural hearing loss, deafferentation

## Abstract

Emerging evidence suggests that cochlear synaptopathy is a common feature of sensorineural hearing loss, but it is not known whether electrophysiological metrics targeting synaptopathy in animals can be applied to a broad range of people, such as those with impaired audiograms. This study investigates the applicability of subcortical electrophysiological measures associated with synaptopathy such as auditory brainstem responses (ABRs) and envelope following responses (EFRs) in older participants with high-frequency sloping audiograms. This is important for the development of reliable and sensitive synaptopathy diagnostics in people with normal or impaired outer-hair-cell function. Broadband click-ABRs at different sound pressure levels and EFRs to amplitude-modulated stimuli were recorded, as well as relative EFR and ABR metrics which reduce individual factors such as head size and noise floor level. Most tested metrics showed significant differences between the groups and did not always follow the trends expected from synaptopathy. Audiometric hearing loss and age-related hearing related deficits interacted to affect the electrophysiological metrics and complicated their interpretation in terms of synaptopathy. This study contributes to a better understanding of how electrophysiological synaptopathy metrics differ in ears with healthy and impaired audiograms, which is an important first step towards unravelling the perceptual consequences of synaptopathy.

## 1. Introduction

The cochlea is a complex structure with many interdependent components that shape how we perceive sound. With age and/or exposure to noise or ototoxic agents, cochlear structures can deteriorate, yielding different degrees and manifestations of sensorineural hearing loss. Outer-hair-cell (OHC) loss is an important contributor to hearing loss and causes a reduced amplification of sensory input that is associated with worsened frequency selectivity and wider auditory filters (Glasberg and Moore, 1986). As OHC loss is associated with elevated pure tone thresholds, this hearing deficit can routinely be quantified using the standard audiogram procedure (Johnson, 1970). However, some aspects of hearing impairment, especially those reported by people with normal-hearing thresholds, are not sufficiently characterised by the audiogram alone (Hind et al., 2011; Kobel et al., 2017; Kumar et al., 2007; Lobarinas et al., 2017). These so-called *suprathreshold* hearing deficits, in the presence of normal sound detection, are a topic of intensive investigation as the underlying cause of these deficits may explain why two individuals with the same audiogram can have very different speech intelligibility scores (Festen and Plomp, 1983).

One possible cause for suprathreshold hearing deficits in the presence of normal hearing sensitivity is *cochlear synaptopathy* or *neuropathy.* Research in mice has shown that an overexposure to noise can lead to a loss of up to 50% of the synapses and cochlear nerve terminals innervating the inner hair cells (IHC) while hearing thresholds are normal (Kujawa and Liberman, 2009). Cochlear synaptopathy is hypothesised to be induced by glutamate excitotoxicity in the post-synaptic terminals of IHCs and the consequences are swelling, bursting and finally the withdrawal of the terminal dendrite (Liberman and Kujawa, 2017). Because each neuron communicates only with one IHC per tonotopic location along the basilar membrane (Stamataki et al., 2006), even a partial deafferentation leads to a loss of information in the chain of information transfer that could eventually lead to neurodegeneration of the spiral ganglion cells (SGC; Liberman and Kujawa, 2017). Low-spontaneous rate (low-SR) fibres with high firing thresholds are relatively more affected by noise exposure (Furman et al., 2013) or aging (Schmiedt et al., 1996) than those with low thresholds and high-spontaneous firing rates (high-SR fibres). Given its expression, synaptopathy is thought to degrade the robust encoding of suprathreshold temporal envelopes (Bharadwaj et al., 2014; Parthasarathy and Kujawa, 2018; Verhulst et al., 2018b), or results in the loss of spectral contrast important for speech-in-noise decoding (Carney, 2018).

In animal studies, it is possible to study and relate histological findings of synaptopathy directly to non-invasive subcortical electrophysiological measures such as the auditory brainstem response (ABR) and the envelope following response (EFR). The most prominent finding is the correlation between the ABR Wave-I amplitude at moderate-to-high sound levels and the number of intact IHC synapses (Furman et al., 2013; Kujawa and Liberman, 2009; Lin et al., 2011). EFRs to modulated pure tones with modulation rates around 0.7-1 kHz can also be a robust and indirect measure for noise or ageing-induced synaptopathy (Parthasarathy and Kujawa, 2018; Shaheen et al., 2015). The relationship between synaptopathy and auditory evoked potentials has opened avenues to diagnose synaptopathy in humans and to study the relationship between synaptopathy and sound perception.

In humans, a stronger reduction of EFR strength with decreasing stimulus modulation depth was found to go along with worse ITD detection thresholds as well as degraded AM detection and poorer selective attention performance in listeners with otherwise normal audiograms (Bharadwaj et al., 2015). Reduced perceptual temporal encoding abilities may thus be diagnosed using this relative EFR slope metric. Second, the ratio between the hair-cell-generated summating potential (SP) and the cochlear-neuron generated action potential (AP or ABR Wave I) was also suggested as a marker of synaptopathy in humans. It predicted the poorer word-recognition-in-noise performance of participants with higher doses of self-reported noise exposure (Liberman et al., 2016). Further studies have related the ABR Wave-I amplitude to the amount of life time noise exposure (Bramhall et al., 2017; Valderrama et al., 2018) and tinnitus (Schaette and McAlpine, 2011) in accordance with noise-induced synaptopathy observations in rodents (Furman et al., 2013; Kujawa and Liberman, 2009; Möhrle et al., 2016). Lastly, Mehraei et al., (2016) argued that increased Wave-V latency for increasing background noise levels may emphasise the contribution of low-SR fibres to the ABR resulting from their high firing thresholds and delayed onset responses (Bourien et al., 2014). Smaller than normal Wave-V latency shifts would hence predict a loss of low-SR fibres.

That synaptopathy is expressed in humans is known from SGC counts performed in post-mortem human temporal bones with normal populations of hair cells, estimating a mean annual loss of up to 100 SGC (Makary et al., 2011; Wu et al., 2018). Work in macaques further suggested that mammals are more resilient to hair cell loss, but show similar vulnerability to cochlear synaptopathy in comparison to most rodent models (Valero et al., 2017). Nevertheless, the degree to which synaptopathy and the use of subcortical measures for diagnostics are transferable to humans is still a topic of debate due to species-specific differences in the physiology of hearing (Hickox et al., 2017; Plack et al., 2016; Prendergast et al., 2017). Furthermore, humans show increased variation in physiological measures compared to animals due to the heterogeneity in tissue-conductance, head size, cognitive abilities, noise exposure across the life span and different genetic factors (Liberman and Kujawa, 2017; Lobarinas et al., 2017; Mitchell et al., 1989; Plack et al., 2016; Trune et al., 1988; Yeend et al., 2017). Especially in the elderly, a relative mix of peripheral and cognitive factors plays a role in speech understanding and suprathreshold auditory temporal processing (Humes, 2013; Humes et al., 2010). This multitude of factors combined with the fact that subcortical EEG measures are only indirect indicators of synaptopathy reduce the translation from animal study findings to individualise diagnostic metrics for cochlear synaptopathy in humans.

Despite a range of positive findings, there is also a considerable body of research that did not find links between sound perception and electrophysiological measures of synaptopathy. For example, a study which investigated the relationship between noise exposure history, suprathreshold functional hearing tests and the ABR Wave-I amplitude did not find any significant relation between the metrics (Fulbright et al., 2017). Guest and colleagues (2017) followed a similar approach and did not find any links between noise exposure history and the ABR Wave-I amplitude, or EFR measures, in young adults with and without tinnitus. Another study which included more than 100 participants with normal audiometric thresholds could not establish any relationship between noise exposure history and the ABR Wave I and the frequency-following response magnitude (Prendergast et al., 2017). From the listed studies, we can either conclude that noise-induced cochlear synaptopathy might not play an important role in young adults with normal audiometric hearing threshold, or that the adopted electrophysiological measures are not sensitive enough to reveal subtle differences in neural fibre populations.

The interpretation of subcortical EEG metrics in terms of synaptopathy is further complicated by the presence of other peripheral contributors to hearing loss, such as OHC deficits. Even though it has been shown that effects of cochlear synaptopathy are more pronounced with advancing age (Fernandez et al., 2015; Parthasarathy and Kujawa, 2018; Sergeyenko et al., 2013), impaired audiograms are also common in the ageing population (ISO, 1990). Aging listeners with impaired audiograms are thus likely to suffer from both OHC deficits *and* synaptopathy, rendering the interpretation of electrophysiological metrics complicated as the metrics can be affected by both deficits. The quantification and isolation of cochlear synaptopathy from other coexisting contributors of hearing loss is therefore still a major unsolved problem in hearing diagnostics (Hickox et al., 2017; Kobel et al., 2017; Plack et al., 2016; Verhulst et al., 2016).

As a first step to disentangle peripheral hearing deficits from a single electrophysiological metric, we investigated whether existing ABR/EFR metrics for synaptopathy diagnosis in a young NH (yNH) group (25±4.1 years) follow the same trends for an older hearing impaired (oHI) group (65±7.9 years) with high-frequency sloping audiograms. The latter group has OHC deficits as verified using the audiogram and distortion-product otoacoustic emission (DPOAE) thresholds (Chen et al., 2008) and is expected to suffer from synaptopathy on the basis of recent studies relating normal ageing to synaptopathy expression (Parthasarathy and Kujawa, 2018; Sergeyenko et al., 2013; Wu et al., 2018). We hypothesise that if synaptopathy drives the considered electrophysiological metrics, the oHI group should perform equally bad or worse than the worst performing yNH participants. However, if the results of the oHI participants do not follow the trends expected from synaptopathy, and show a relationship to hearing sensitivity differences within the older group, the considered metric is likely impacted by both OHC and synaptopathy and may consequently not be a robust marker of synaptopathy in listeners with impaired audiograms. This study investigates which differences in markers of synaptopathy developed for NH listeners can realistically be expected in two extreme participant groups: a young reference group (yNH) and a representative clinical population with mild sensorineural hearing loss (oHI). The study outcomes can help to restrain the large parameter space of potentially appropriate diagnostic metrics for synaptopathy to identify measures that can quantify synaptopathy in the presence of normal *or* abnormal OHC function.

### 1.1 Considered ABR/EFR metrics and expected outcomes

We report individual differences in electrophysiological response behaviour for multiple stimulus parameters such as bandwidth, modulation frequency and depth as well as sound pressure level (SPL) to provide a comprehensive view of the applicability of subcortical EEG metrics in the two listener groups. We incorporate a relative metric design to account for inter-individual differences and further reduce measurement uncertainty by combining different measures. In our analysis, we consider relationships of the metrics to objective physiological markers of peripheral hearing and other electrophysiological markers. We do not specifically consider age as an explanatory variable as it can be linked to both synaptopathy (e.g. Parthasarathy and Kujawa, 2018) and OHC deficits (e.g. Wu et al., 2018). We investigate the following hypotheses which are motivated by extending the NH synaptopathy results to an older HI group with high-frequency hearing loss and a suspected high degree of synaptopathy.

#### 1) Reduced EFR amplitudes

On the basis of animal-synaptopathy findings (Parthasarathy and Kujawa, 2018; Shaheen et al., 2015) and reduced temporal envelop encoding ability (Bharadwaj et al., 2014) we expect that the EFR strength is reduced in the oHI group. Because biophysical EFR-model predictions support the idea that OHC loss does not strongly influence the EFR metric (Verhulst et al., 2018a, 2018b, 2016), we predict that reduced EFR amplitudes in the oHI group predominantly reflect their individual degree of synaptopathy. In line with this, individual differences in oHI-EFRs should not relate to hearing threshold differences in this group.

#### 2) Steeper EFR slope metric

If the EFR strength reflects the temporal coding ability of the brainstem, the EFR amplitude slope as a function of modulation depth reduction should be steeper in the oHI group. If this is not the case, the EFR slope metric might not exclusively be sensitive to synaptopathy and reflect a contribution of OHC deficits as well.

#### 3) Lower ABR amplitudes for equal SPL at suprathreshold levels

The ABR Wave I should be reduced in the oHI group if synaptopathy is driving this metric (e.g. Kujawa and Liberman, 2009). We also expect reduced amplitudes for the Wave V if central gain compensation does not play a role. Because central gain compensation (i.e., normal Wave V in the presence of a reduced Wave I) was mostly observed in young, but not older, animals with synaptopathy (Möhrle et al., 2016), we expect that the ABR Wave-V amplitudes of our advanced-age oHI group relate well to how synaptopathy affects the ABR Wave-I.

#### 4) Shallower ABR amplitude slopes

If synaptopathy is the driving force behind the ABR amplitude as a function of stimulus level increases (i.e., the ABR amplitude slope), we expect a shallower slope for the oHI group. However, a potential loss of cochlear compression due to OHC deficits would yield *steeper* ABR amplitude slopes. OHC deficits are also associated with longer low-level ABR latencies (Lewis et al., 2015) and steeper ABR latency slopes (Gorga et al., 1985; Verhulst et al., 2016) based on relatively stronger contributions of the intact apical portion of the basilar membrane. If synaptopathy drives the ABR amplitude metric, we expect to see shallower ABR slopes which do not relate to the ABR latency slope, expected to reflect OHC loss.

#### 5) Relationship between EFR and ABR metrics

Lastly, if the ABR amplitude slope and EFR slope both represent aspects of synaptopathy, we expect a positive correlation between these metrics under the assumption that the neuronal population generating the EFR can be seen as a subgroup of the fibres responding to the click-ABR. Likewise, we expect to see a relationship between individual EFR magnitudes and the EFR slope as a function of stimulus modulation depth, and of the EFR magnitude with the ABR amplitude slope.

## 2. Materials and Methods

### 2.1 Participants

The study included 23 young normal-hearing (yNH) participants between 14 and 32 years of age (M_age_ = 25 years; SD_age_ = 4.1, 14 females) and 23 older hearing-impaired participants (oHI) aged between 48 and 77 (M_age_ = 65 years; SD_age_ = 7.9, 11 females). All yNH participants had normal pure tone thresholds (≤ 25 dB HL) assessed with a clinical audiometer (Auritec AT900, Hamburg,Germany) using a standard procedure for frequencies between 0.125 and 8 kHz. The oHI participants had high-frequency sloping audiograms with hearing losses up to 45 dB HL at 4 kHz as shown in the red/orange traces in Figure 1. Given the evidence that synaptopathy sets in before OHC loss can be detected (Fernandez et al., 2015; Parthasarathy and Kujawa, 2018; Sergeyenko et al., 2013; Wu et al., 2018) the oHI group has expected sensory as well as synaptic hearing damage. To assess OHC integrity and cochlear function, audiometric thresholds were complemented with DPOAE thresholds measured at 4 kHz. The DPOAE thresholds roughly correlated with the pure tone detection threshold measured at the same frequency using a two-alternative forced-choice adaptive tracking procedure and insert earphones (ER-2 with ER10-B, Etymotic Research, Elk Grove Village, IL, USA). Further details on the DPOAE threshold procedure can be found in Verhulst et al. (2016). Participants were informed about the experimental procedures according to the ethical guidelines at the University of Oldenburg. Written informed consent was obtained and participants were paid for their participation.

**Fig. 1:**
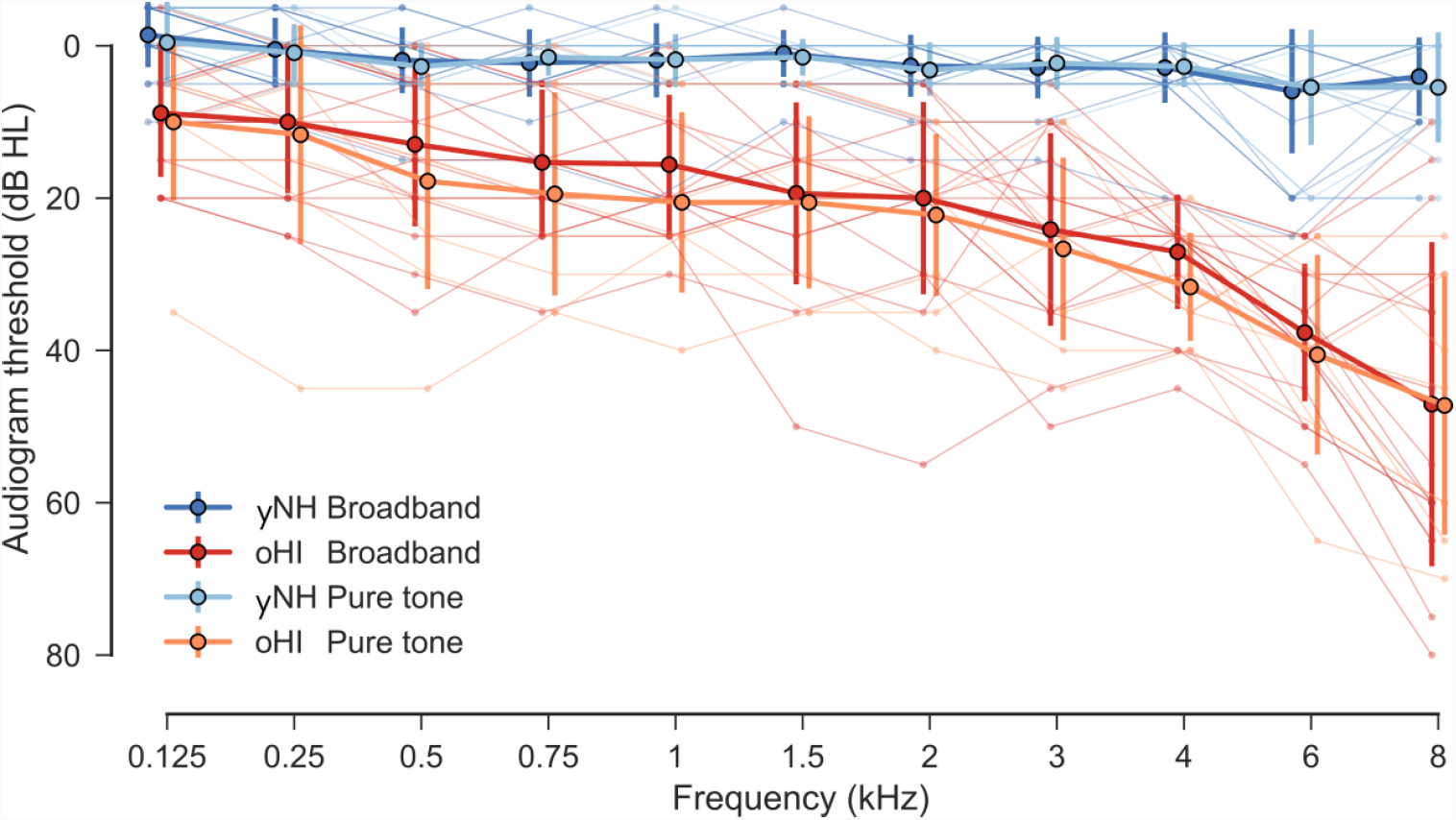
Pure tone hearing thresholds (dB HL) at octave frequencies between 0.125 and 8 kHz. Groups are split according to the degree of hearing loss (young normal-hearing - yNH and old hearing-impaired - oHI) and main EFR stimulus conditions (broadband and pure tone). Thick lines indicate the mean and error bars display the standard deviations across groups. The thin lines represent individual audiogram traces.

### 2.2. Stimuli

Each ABR epoch had a duration of 30 ms and consisted of a 80-µs condensation click followed by silence. After each epoch, a short uniformly distributed random silence jitter (> 0 and < 3 ms; mean = 1.5 ms) was added. 7000 epochs were presented at a rate of 33.3 Hz for all four tested conditions: 70, 80, 90 and 100 dB peak-equivalent sound pressure level (peSPL).

The amplitude-modulated EFR stimuli consisted of two main conditions. The broadband (BB) condition was designed to achieve a maximally broad excitation on the basilar membrane while the pure tone (PT) was expected to maximise individual differences in the 4-kHz range, where the DPOAE threshold was measured. Several other EFR conditions were collected from a subset of participants (see Table 1). The stimuli varied in bandwidth, SPL, modulation depth (MD) and modulation frequency (f_m_).

**Tab. 1:** Overview of the stimulus conditions and their parameters. Sound pressure level is given as dB SPL and the number of participants (n) is shown per condition. The modulation depth (MD) is in dB relative to 100% modulation.

|  | Broadband 120 Hz | Pure tone 120 Hz | Narrowband 120 Hz | Broadband 480 Hz |
| --- | --- | --- | --- | --- |
| yNH | 75 dB; n = 21;<br>MD = 0, -4, -8 | 70 dB; n = 11;<br>MD = 0, -4, -8 | 75 dB; n = 21; MD = 0<br>70 dB; n = 11; MD = 0 | 75 dB; n = 21; MD = 0 |
| oHI | 75 dB; n = 17;<br>MD = 0, -4, -8 | 70 dB; n = 9;<br>MD = 0, -4, -8 | 75 dB; n = 14; MD = 0<br>70 dB; n = 9; MD = 0 | 75 dB; n = 17; MD = 0 |

The BB stimuli consisted of a 75-dB-SPL white noise carrier that was amplitude-modulated with a modulation frequency of 120 Hz. The 4-kHz PT stimuli were calibrated to 70 dB SPL. For both stimulus types, three different modulation depths 0, −4, −8 dB (equivalent to 100, 63 and 40 % depth) were used yielding a total of six main conditions. All other conditions were only tested at 100% modulation depth. The narrowband (NB) stimuli had the same white noise carrier as the broadband stimuli but were band-limited to one octave centred around 4 kHz. This stimulus was designed to achieve good signal strength while retaining frequency specificity around the centre frequency. The narrowband stimuli were tested at 75 and 70 dB SPL to allow for a comparison to the broadband and the pure tone data, and to assess the influence of SPL on the EFR. The ‘broadband 480’ condition only differed from the broadband condition by its modulation frequency of 480 Hz. This higher modulation frequency was included to target more peripheral generators than the brainstem (Purcell et al., 2004) and might be more directly related to AN processing.

Each broadband stimulus lasted 600 ms followed by a uniformly distributed random silence jitter (>90 and <110 ms; mean = 100 ms). All narrowband and pure-tone stimuli were ramped using a 5% tapered-cosine window and were repeated 800 times, whereas all broadband stimuli were repeated only 600 times because they elicited more robust EFRs. Both polarities (50% each) were presented. Due to time restrictions, not all participants were tested in all possible conditions. Three yNH and nine oHI participants took part in both main conditions (BB, PT), all other participants were only exposed to either the BB or PT condition. EEG recording took place in a double-walled electrically shielded measurement booth. Participants sat comfortably in a reclining chair while watching a silent movie. All stimuli were presented monaurally (better ear based on the audiogram) using foam tips coupled to magnetically-shielded ER-2 insert earphones (Etymotic Research, Elk Grove Village, IL, USA) which were connected to a TDT-HB7 headphone driver (Tucker-Davis, Alachua, FL, USA) and a Fireface UCX sound card (RME, Haimhausen, Germany). All stimuli were generated in MATLAB at a sampling rate of 48 kHz and calibrated using an oscilloscope (for ABR only), B&K type 4157 ear simulator and sound level meter type 2610 (Brüel & Kjær, Nærum, Denmark). EEGs were recorded using a 32-channel EEG amplifier and cap (Biosemi, Amsterdam, Netherlands) with a sampling rate of 16384 Hz and 24- bit AD conversion. Common mode sense active and driven right leg passive electrode (CMS/DRL) were placed near the vertex of the participant. The data were re-referenced to the offline averaged earlobe electrodes. The vertex electrode (Cz), yielding the best signal strength, was used for all further analyses. Electrode offsets (DC values of the common mode signal) were kept below 20 mV.

### 2.3. Data Processing and Analysis

The ABR data were filtered using a finite impulse response filter (FIR) with a forward-backward procedure to avoid phase shifts. The data were first high-pass filtered at 200 Hz after which they were low-pass filtered at 1500 Hz. After filtering, the data were epoched between −5 and 20 ms around the stimulus onset, baseline corrected and averaged. ABR latencies and peak amplitudes of Wave I and V were extracted using the auditory wave analysis tool for Python developed by B. Buran (see https://github.com/bburan/abr for details on the procedure). All reported latencies were compensated for by the fixed recording delay of the sound delivery system (1.16 ms).

All EFR data were pre-processed using the Python programming language (version 2.7.10 | Anaconda 2.3.0 (64-bit), www.python.org) and MNE-Python (version 0.9.0) (Gramfort et al., 2014, 2013). An EOG channel was constructed from the electrodes Fp1 and Fp2 to facilitate eye movement artefact detection. To improve the Independent Component Analysis (ICA) results, the data were filtered (1 to 40 Hz) using an infinite impulse response (IIR) Butterworth filter of 4_th_ order. They were epoched to one second long chunks using an epoch rejection threshold of 150 µV. The implemented fast ICA algorithm was applied (max. 300 iterations) and eye movement related artefacts were determined (Debener et al., 2010). The ICA weights were then applied to the original unfiltered data to remove artefacts.

The eye-artefact-free data were high-pass filtered at 60 Hz and then low-pass filtered at 650 Hz using a 4_th_ order IIR Butterworth filter. A zero-phase shift was achieved by applying a forward-backward filter procedure. Bad channels were removed based on recording notes and visual inspection. Data were epoched from −0.01 to 0.6 sec. around trigger onset. The 10 ms before trigger onset were used for baseline correction and discarded afterwards. Epochs with amplitudes exceeding a 100 μV threshold were removed. Each epoch was transformed to the frequency domain using Matlab’s Fast Fourier Transform (FFT) function. To estimate the frequency dependent noise floor and to get a better estimate of the data distribution, a bootstrap procedure was applied (Zhu et al., 2013). First, a magnitude spectrum estimate of the neural responses for each condition and participant was computed by averaging randomly drawn epochs. The random draws (with replacement) equalled the number of epochs left after artefact rejection in each condition. This step was repeated 200 times resulting in an estimated magnitude spectrum distribution. The average spectrum of this approximately Gaussian distributed measure was used as the estimate of the individual participant’s magnitude spectrum of the response per condition. The standard deviation of the 200 estimates was used as an estimator of the variability. The spectral magnitude of the noise floor was calculated using a similar approach, except for that the number of frequency estimates was increased to 1000 and that the phase of half of the randomly drawn epochs were flipped. This method cancels out the constant time-locked signal (i.e., the EFR) in the recording and only preserves the non-stationary noise that has a characteristic shape proportional to 1/f (Voytek et al., 2015). The estimated noise floor and the response estimate were then transformed to a logarithmic dB scale using 20*og_10_ (). The reference reflects the amplitude of a pure tone with a root-mean-square value of one. Lastly, the estimated noise floor was subtracted from the absolute EFR magnitude to yield a signal-to-noise ratio measure (EFR SNR) which was used in all further EFR analyses and is referred to as EFR or EFR magnitude. Responses were considered as significant above the noise floor if their magnitude at the modulation frequency exceeded the 1000 computed noise floor estimates in more than 95% of all cases. The EFR normalization to the noise floor allows for a better comparison of EFR SNRs between individuals as it takes into account individual differences in background noise floor level. The EFR normalization is also particularly important when comparing EFRs across different modulation frequencies as both the modulation transfer function (Purcell et al., 2004; Tichko and Skoe, 2017) and the background noise levels are frequency dependent.

The assumptions for the performed statistical inference tests were tested using the ‘sciPy’ python package for scientific computing (Millman and Aivazis, 2011; Oliphant, 2007). The assumption of normal-distribution was tested using the Shapiro-Wilk-Test. The equal-variance assumption was tested using the Leven-Test. If assumptions were satisfied dependent/independent t-tests were used to test differences between two samples. If the normal-distribution assumption was not met for two independent samples, the non-parametric Mann-Whitney U-Test (U) was applied. If only the equal-variance assumption was violated, Welch’s t-test was performed. If two dependent samples violated the normal-distribution assumption, the Wilcoxen signed-rank test (W) was applied. All correlations reported refer to the Pearson correlation coefficient (r) if both variables were normally-distributed, otherwise Spearman’s rank correlation coefficient (ρ) was used.

All analyses of variance (ANOVA) and post-hoc analyses were done in the R programming environment (R Core Team, 2017) using the ‘nlme’ (Pinheiro et al., 2017) and ‘lsmeans’ (Russell, 2016) packages. All reported p-values for multiple comparisons were Bonferroni adjusted to control for the family-wise error rate and can be directly compared to the applied significance level of α = 0.05.

## 3. Results

The EFR magnitudes for the main BB and PT stimulus conditions and three modulation depths are illustrated in Figure 2 for the different participant groups. A two-factor (2×3) mixed design ANOVA comparing all data for the BB-EFR magnitudes indicated a significant main effect of the group (F(1,36) = 61.57; p < 0.0001; η2 = 0.63) and modulation depth (F(2,72) = 63.24; p < 0.0001; η_^2^_ = 0.62) but no significant interaction. A post-hoc analysis of pairwise comparisons (15 comparisons) showed significantly lower EFR magnitudes for the oHI participants in all three modulation depth conditions. As expected, decreasing the modulation depth resulted in smaller EFRs. EFR magnitudes were significantly smaller between the 0 dB and −8 dB conditions as well as for the −4 dB and −8 dB contrast in both groups. No significance was indicated between the 0 dB and −4 dB conditions in neither of the groups. The PT-EFRs showed similar outcomes with a significant main effect of group (F(1,18) = 26.34; p < 0.0001; η_^2^_ = 0.59) and modulation depth (F(2,36) = 6.08; p = 0.0053; η_^2^_ = 0.23). The interaction did not reach significance. The post-hoc tests (15 comparisons) only showed significant differences between the groups for the 0 and −4 dB modulation depth conditions, and between the 0 and −8 dB conditions within the yNH group. When only considering the EFRs that exceeded the noise floor significantly in the analysis (see methods), the reduction of EFR magnitudes with decreasing modulation depth was strongly diminished for the PT and BB conditions in the oHI group and for the PT condition in the yNH group. The percentage of participants with EFR magnitudes above the noise floor is shown in Table 2 for the BB and PT conditions.

**Fig. 2:**
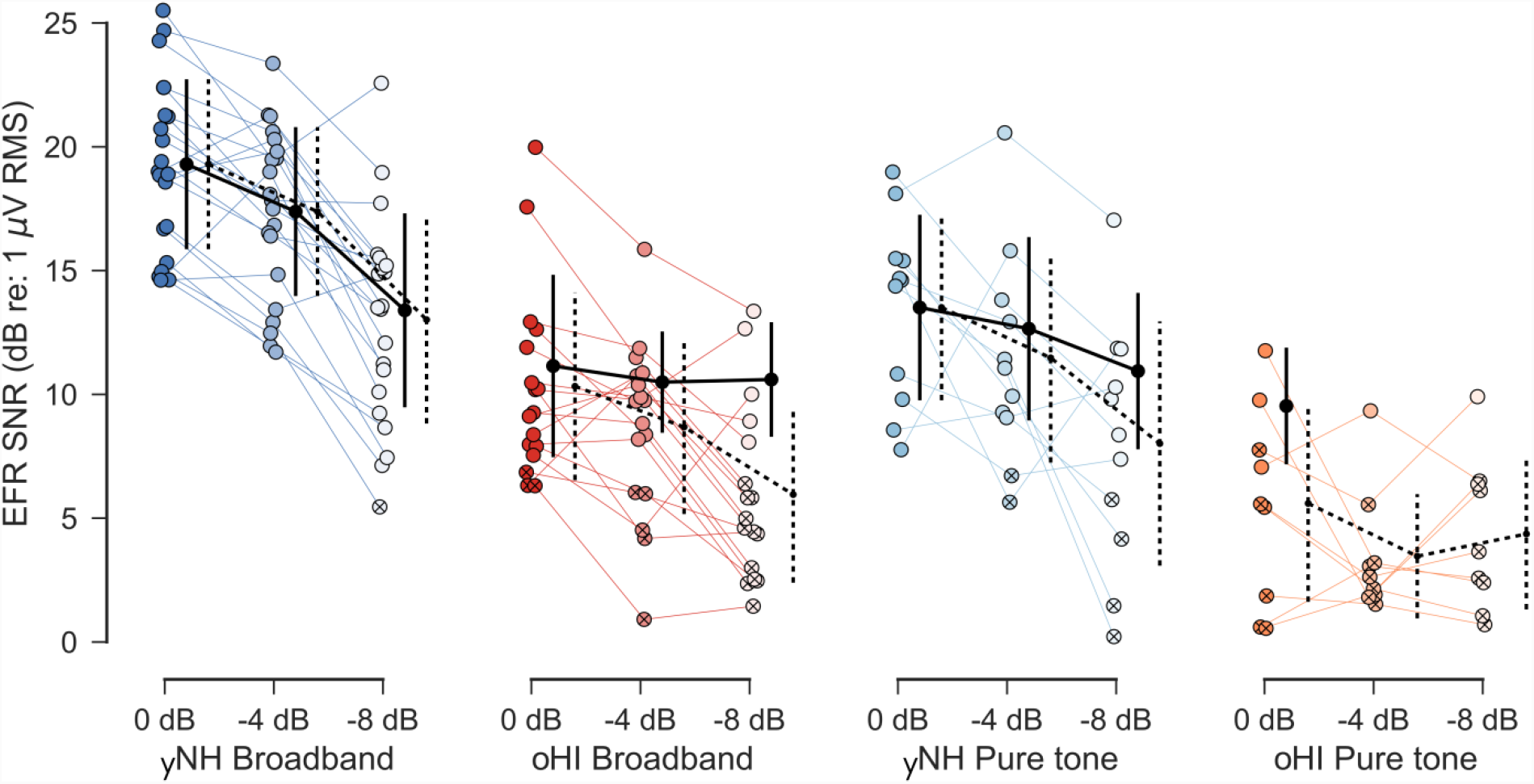
Individual traces of the bootstrapped EFR SNR magnitudes for both main stimulus conditions (BB, PT) and participant groups (yNH, oHI) over all three modulation depth conditions (0, −4, −8 dB). Crosses indicate values at noise floor level. Error bars show mean and standard deviation of significant data (solid) and all data (dashed) across groups.

**Tab. 2:**
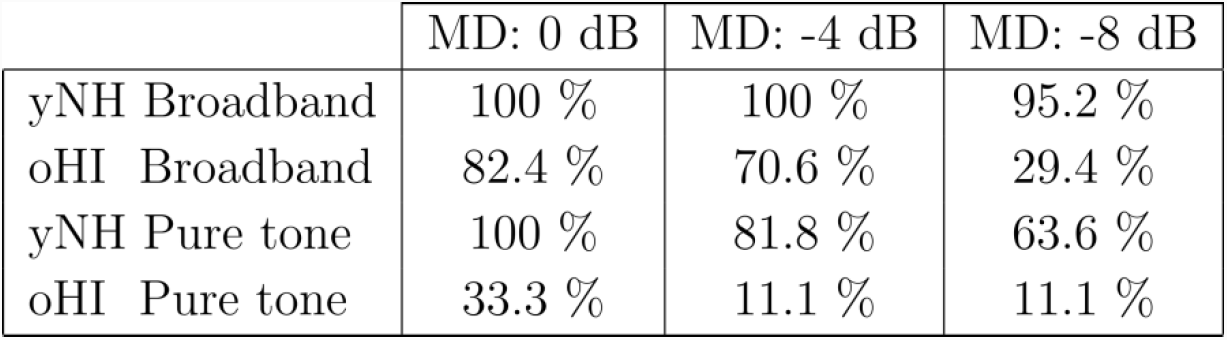
Percentage of participants per carrier (BB,PT), group (yNH/oHI) and modulation depth condition (0,-4,-8 dB re:100%) whose raw estimated EFR response at the modulation frequency (120 Hz) exceeded the 1000 individually estimated noise floor values more than 95% of the time.

|  | MD: 0 dB | MD: -4 dB | MD: -8 dB |
| --- | --- | --- | --- |
| yNH Broadband | 100 % | 100 % | 95.2 % |
| oHI Broadband | 82.4 % | 70.6 % | 29.4 % |
| yNH Pure tone | 100 % | 81.8 % | 63.6 % |
| oHI Pure tone | 33.3 % | 11.1 % | 11.1 % |

The increase of non-significant EFRs for smaller modulation depths, especially prominent in oHI participants, underlines the increased difficulty for the brainstem to encode envelope information robustly. The influence of the carrier type (BB vs. PT) was investigated using a two-factor (2×3) mixed design ANOVA. The carrier type shows a significant main effect for the yNH (F(1,30) = 21.01; p = 0.0001; η2 = 0.41) and oHI participants (F(1,24) = 10.56; p = 0.0034; η2 = 0.31). The BB carrier shows larger EFR values, as confirmed in a post-hoc analysis (15 comparisons) for all MD conditions in the yNH group (0.0050 ≤ p ≤ 0.0268) and for the 0 and −4 dB conditions (0.0207 ≤ p ≤ 0.0477) in the oHI group.

To reduce the potential influence of confounding factors such as head size, skull thickness and gender on the EFR magnitudes, a relative self-normalizing slope metric to quantify suprathreshold coding fidelity was suggested (Bharadwaj et al., 2014). The data points (m ≈ 1, 0.63, 0.4) corresponding to 20 log10(m) = 0, −4, −8 dB modulation depth were used to fit a straight line through the EFR magnitudes as a function of the stimulus modulation depth. All data points, including those below the noise floor, were used to calculate the slopes. EFR slopes are shown in Figure 3 and the yNH values corroborate the slopes reported in other studies (Bharadwaj et al., 2015; Guest et al., 2017). The PT-EFR slopes showed a greater variance than the BB-EFR slopes in the yNH group and independent two-sided t-tests comparing yNH and oHI participants in the BB (t(36) = −1.90; p = 0.0660) and PT conditions (t(18) = −1.63; p = 0.1209) did not indicate significant differences. This result was unexpected given that shallower EFR slopes are associated with better temporal envelope coding ability (Bharadwaj et al., 2015) and as the oHI group is expected to suffer from synaptopathy, they should have steeper-than-normal slopes. Opposite to what was expected from the synaptopathy-hypothesis, oHI participants had shallower mean slopes than the yNH group, and some participants even showed positive slopes. To better understand the observed slopes, we investigated how the EFR slopes are linked to the EFR magnitudes (see Figure 4). The yNH-EFR slopes only showed a significant correlation to the −8-dB MD EFR condition (BB: r = −0.52, p = 0.0157; PT: r = −0.76, p = 0.0062) reflecting that the EFR slope is mostly determined by the degree of temporal coding at lower modulation depths. On the other hand, the oHI slopes showed an inverse relationship as the HI-EFR slopes correlated strongest with the 0-dB MD EFR condition (BB: ρ = 0.47, p = 0.0551; PT: r = 0.76, p = 0.0169). Their slope is therefore mainly determined by the degree of temporal coding at higher modulation depths.

**Fig. 3:**
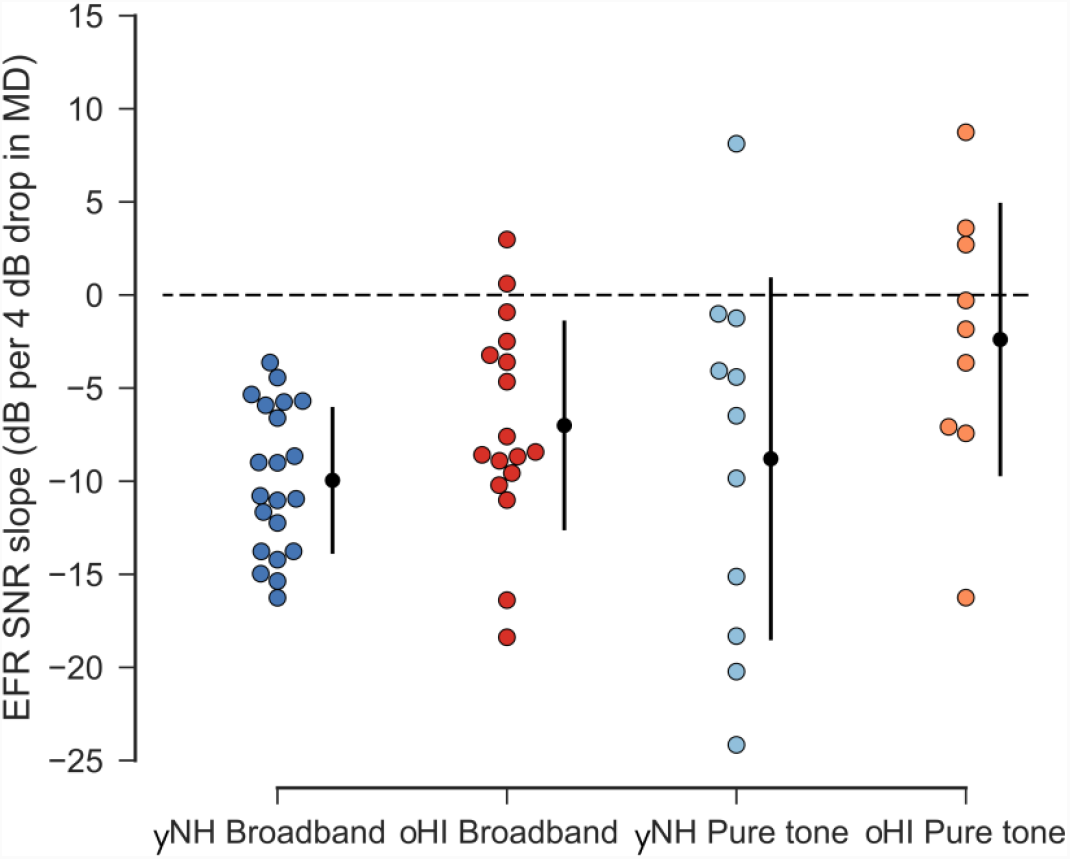
EFR SNR slopes (slope of a straight line fitted through all three modulation depth conditions) for both stimulus conditions (BB, PT) and groups (yNH, oHI). More negative values indicate a steeper EFR SNR reduction with decreasing modulation depth.

**Fig. 4:**
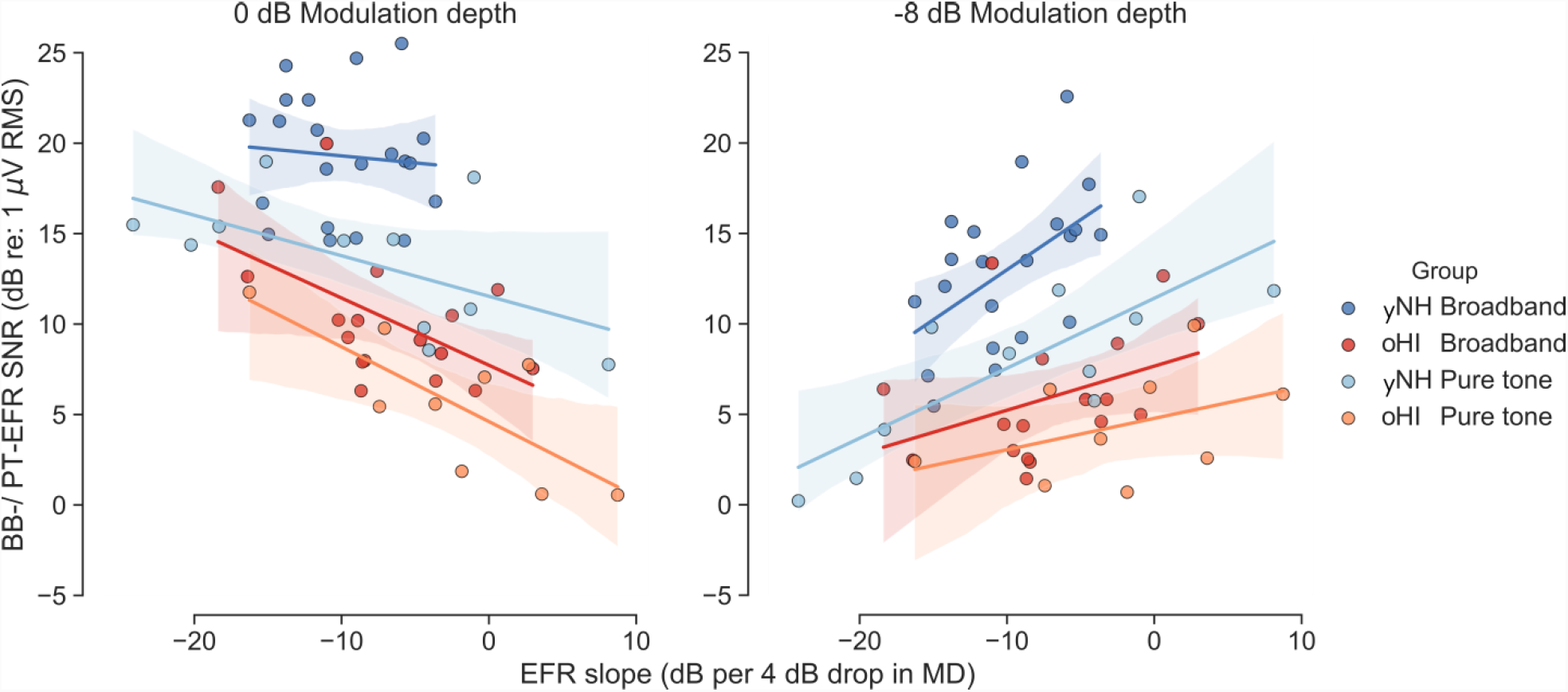
Regression plot of the EFR SNR slopes (slope of a straight line fitted through all three modulation depth conditions) with the 0 dB-MD EFR SNR magnitudes (A) and −8 dB-MD EFR SNR magnitudes (B) for both main stimulus conditions (BB, PT) and participant groups (yNH, oHI). The shaded areas display the 95% confidence interval of the regression fit.

With one exception, we did not find significant correlations between the EFR magnitudes in the different conditions and the 4-kHz audiogram or 4-kHz DPOAE thresholds within the yNH or oHI group. Given that synaptopathy does not affect hearing thresholds (Kujawa and Liberman, 2009), these results fall within the expectation that suprathreshold temporal envelope encoding is independent of hearing sensitivity. However, at 4 kHz, we did see a single positive correlation between the PT-EFR (100% MD) and the audiogram threshold within the oHI group (r = 0.68, p = 0.0430, N = 9). Of all oHI participants, the largest EFR magnitudes were found for individuals with audiogram thresholds > 30 dB HL (i.e. greater hearing loss). This result might be explained by a mechanism in which greater degrees of OHC loss result in a linearization of cochlear processing, which could counteract the EFR reduction associated with synaptopathy. Significant EFR magnitudes were recorded for the majority of oHI participants for 100% modulated stimuli, suggesting that the EFR metric itself might be robust for usage in listeners with sloping high-frequency audiograms. However, as the PT-EFRs were only significant above the noise floor for individuals with thresholds > 30 dB HL, OHC deficits might interact with synaptopathy to affect the EFR magnitude (at least in the PT condition). The only significant link between the hearing sensitivity measures (4-kHz audiogram/DPOAE thresholds) and the EFR slope metric was found in the oHI-PT group (r = 0.86, p = 0.0032) which can be explained by how the slope links to the EFR magnitudes (see above).

### 3.1. Effect of bandwidth and sound pressure level

A stimulus bandwidth change from BB to NB (t(20) = 5.85; p < 0.0001) and again from NB to PT (t(10) = 4.76; p = 0.0009) reduced the yNH-EFRs significantly (left panel of Figure 5A). A significant effect of bandwidth on the oHI-EFRs (right panel of Figure 5A) was not found. On a group level and using an independent t-test, a 5-dB stimulus level difference reached significance (t(30) = −2.20; p = 0.0361), showing a larger EFR mean for the 70-dB-SPL than for the 75-dB-SPL NB condition. This level effect was still present when only including participants who participated in both conditions (t(8) = −2.53, p = 0.0354). For the oHI participants, this difference only reached significance on a group level when considering all data points (t(21) = 2.60, p = 0.0168). The oHI participants showed significantly smaller EFRs than the yNH participants in both NB conditions (75 dB: t(33) = 4.45; p = 0.0001 and 70 dB: t(18) = 7.35; p < 0.0001).

**Fig. 5:**
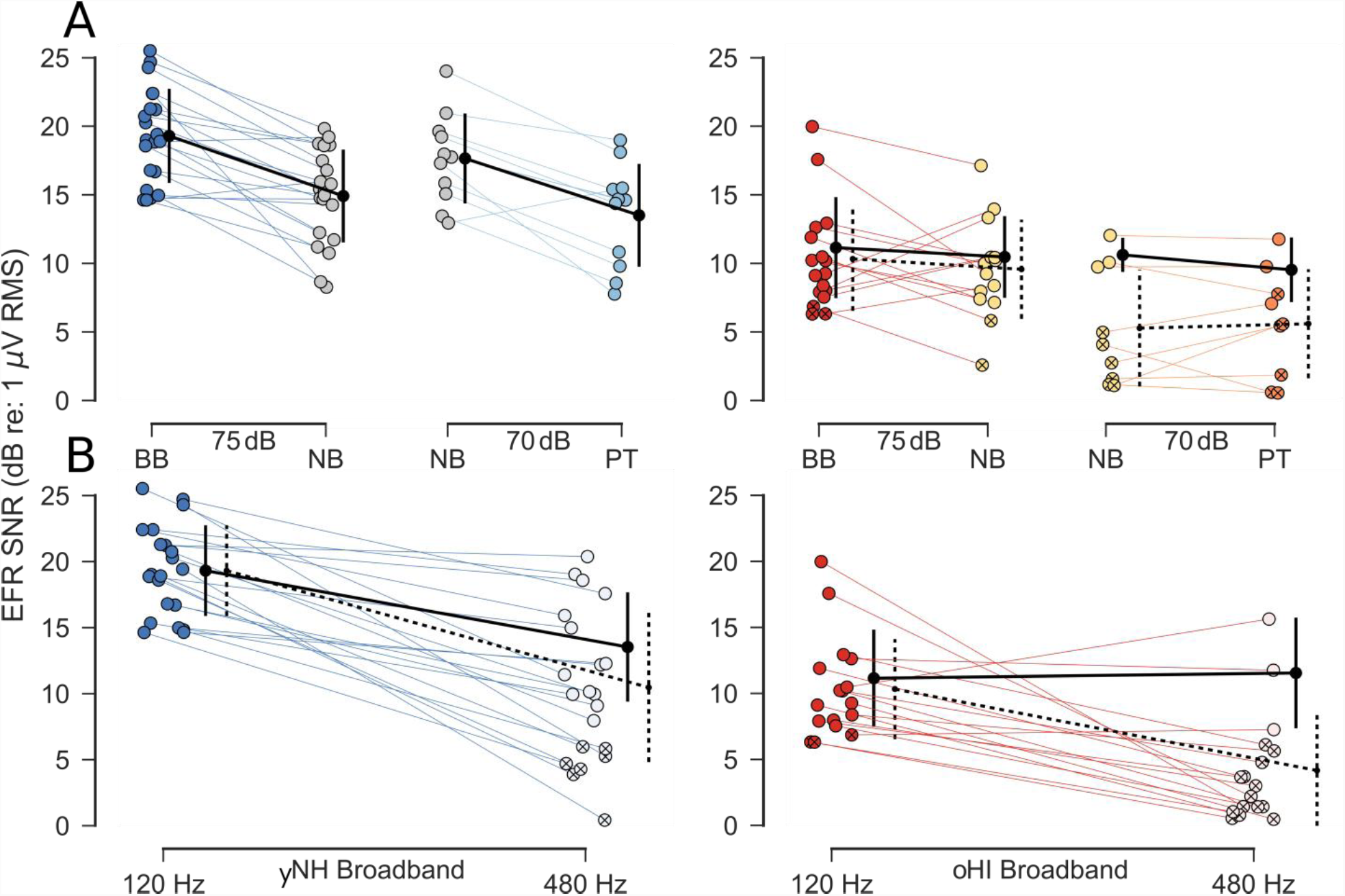
Influence of different stimulus parameters on the EFR SNR response in the 0 dB modulation depth condition for a subset of participants in both groups (yNH, oHI). A) Effects of bandwidth and SPL. Single traces (where available) represent each participant’s response over different bandwidth conditions (BB and NB at 75 dB; NB and 4-kHz PT at 70 dB). B) Effects of modulation frequency. Single traces represent each participant’s response over different modulation frequencies (120, 480 Hz) in the BB condition. Crossed markers indicate values at noise floor level. Error bars show mean and standard deviation of significant data points (solid) and all data points (dashed) across participants in a group.

### 3.2. Effect of modulation frequency

When comparing EFR magnitudes to the modulation frequencies of 120 and 480 Hz (Figure 5B), the yNH participants showed a significant homogenous decrease with increasing modulation frequency (t(20) = 7.40; p < 0.0001), even when only considering significant responses (t(13) = 5.80; p = 0.0001). The correlation between the two EFR conditions showed a positive trend but did not reach significance (r = 0.35; p = 0.12, N = 21). For oHI participants, where the majority of 480-Hz EFRs were below the noise floor level, the modulation frequency increase also reduced the EFR magnitude significantly (W = 7.0; p = 0.001). An independent t-test between groups for the 480-Hz conditions revealed that yNH participants had significantly larger EFRs (U = 63.0, p = 0.0004) than the oHI group.

### 3.3. Auditory Brainstem Responses

The ABR traces were analysed for Wave I and V, and their respective amplitudes and latencies are shown in Figure 6. The insets show raw grand-average traces of the yNH (blue) and oHI (red) participants for all four level conditions. A two-factor mixed design ANOVA for Wave I indicated significantly larger amplitudes for yNH participants (F(1,44) = 9.67; p = 0.0033; η_^2^_= 0.18) but a post-hoc analysis (28 comparisons) showed that this was only the case for the 90 dB peSPL condition (t-ration = −4.106; p = 0.0048). No main effect was found for the increase in peSPL. The interaction shows significance (F(3,132) = 2.88; p = 0.0381; η_^2^_ = 0.06) resembling the large variability in the Wave-I data (panel 6A). The average amplitudes for the oHI participants did not show a consistent increase with peSPL. The same analysis for the Wave-I latency showed a significant main effect of peSPL (F(3, 132) = 83.70; p < 0.0001; η_^2^_ = 0.64) which was similar for both groups and showed no interaction.

**Fig. 6:**
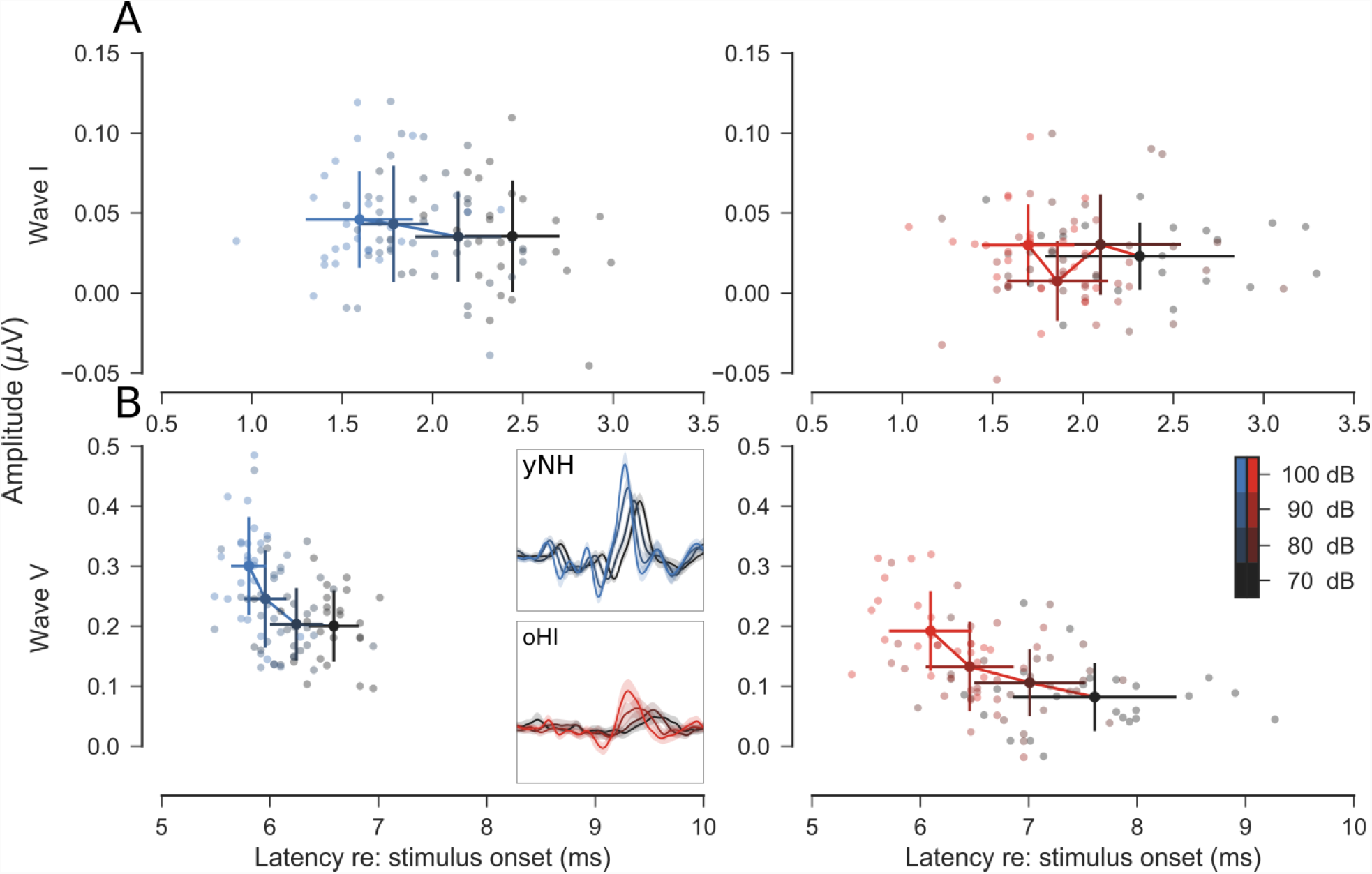
ABR latency vs. amplitude scatter plots for all yNH and oHI participants over the four peSPL levels for Wave I (A) and Wave V (B). Error bars display means and standard deviations across all participants in a group. Insets display the raw grand average ABR traces with 95% confidence intervals (filtered between 200 and 1500 Hz) for both participant groups. PeSPLs are indicated using colours ranging from black (70 dB) to blue/red (100 dB) where lighter colours represent higher levels (see colour bar).

The analysis for Wave V showed more reliable results compared to the Wave-I analysis. An ANOVA for the Wave-V amplitudes indicated significant effects of group and peSPL. The yNH participants showed larger amplitudes (F(1,44) = 37.37; p < 0.001; η_^2^_ = 0. 46) than the oHI participants in all four peSPL conditions as revealed by a post-hoc analysis (28 comparisons) as depicted in panel 7A. Wave-V amplitudes increased in both groups with increasing peSPL (F(3,132) = 81.02; p < 0.0001; η_^2^_ = 0.64) and post-hoc pairwise comparisons (28 comparisons) confirmed this for most conditions. Exceptions that did not reach significance were the 70-vs.-80 dB-peSPL contrast in the yNH group, and the 70-vs.-80 and 80-vs.-90 dB-peSPL contrast in oHI participants. No significant interaction between group and peSPL was found. In contrast to the Wave-I latencies, Wave-V latencies were significantly shorter in yNH participants (F(1,44) = 41.85; p < 0.0001; η_^2^_ = 0.49), except for the 100 dB-peSPL condition (t-ratio = 2.41, p = 0.56). The latencies in both groups converged to each other for higher peSPLs and showed a clearly visible reduction in variance with increasing peSPL for the oHI participants (see panel 7B). Only at the 100 dB peSPL condition and only for the oHI participants, did the ABR amplitudes and latencies show a significant correlation (r = −0.49, p = 0.0176, N = 23), suggesting that at higher peSPL, the latency as well as the amplitude reflect the degree of OHC loss. The peSPL had a significant main effect (F(3, 132) = 169.16; p < 0.0001; η_^2^_ = 0.74) on the yNH Wave-V latency, showing a systematic decrease with increasing peSPL. Post-hoc pairwise comparisons (28 comparisons) for all but the yNH 90-vs-100 dB-peSPL condition (t-ratio = −1.99, p = 1.00) reached significance, corroborating earlier reports (Lewis et al., 2015). All contrasts for the oHI participants reached significance, indicating a stronger influence of peSPL on the oHI latencies at higher peSPLs compared to the yNH participants, which is supported by a significant interaction effect (F(3, 132) = 16.81; p < 0.0001; η2 = 0.07). This again points to OHC loss as being the driving influence behind this metric. This is further supported by the significant correlations of the ABR Wave-V amplitude (70 dB: ρ = −0.50, p = 0.0147; 80 dB: ρ = −0.54, p = 0.0074; 100 dB: ρ = −0.55, p = 0.0065, N = 23) and the Wave-V latency (90 dB: ρ = 0.41, p = 0.0499; 100 dB: ρ = 0.52, p = 0.0103, N = 23) with the 4-kHz audiometric threshold only found in the oHI group. Smaller amplitudes and longer latencies go along with higher audiometric thresholds. The 4-kHz DPOAE thresholds which were reliably extracted from the recordings also showed significant correlations with the ABR Wave-V amplitudes in all four tested peSPL conditions (0.0025 ≤ p ≤ 0.0083, N = 13), but again only in the oHI group.

Similar to the EFR data, relative slope measures were computed from the Wave V ABR data to reduce the effects of confounding factors on the absolute amplitudes. Such relative changes in ABR characteristics have successfully been linked to cochlear synaptopathy in the past (Mehraei et al., 2016). A straight line was fitted for each subject through the four data points representing the amplitude and latency information. Figure 8 depicts the resulting ABR latency (panel A) and ABR amplitude slope values (panel B) as a function of increasing peSPL for both groups. The yNH participants showed very homogenous responses in both measures in comparison to the oHI participants. yNH participants showed significantly shallower positive amplitude slopes (U = 87.0; p = 0.0001). Due to the violation of the equal variance assumption, Welch’s t-test for independent samples was used to assess the latency slope differences between groups. The yNH group showed shallower negative latency slopes (t(24.95) = 4.91; p < 0.0001) than the oHI group. The steeper oHI amplitude slopes did not follow the trend expected from synaptopathy which predicts shallower amplitude growth for participants with synaptopathy (Furman et al., 2013). The observed steeper negative oHI latency slopes fall in line with how sloping high-frequency audiograms are expected to affect the ABR latency slope (Gorga et al., 1985). At the same time, the ABR slope metrics did not link to any of the recorded hearing sensitivity measures (4-kHz audiometric threshold or 4-kHz DPOAE thresholds).

**Fig. 7:**
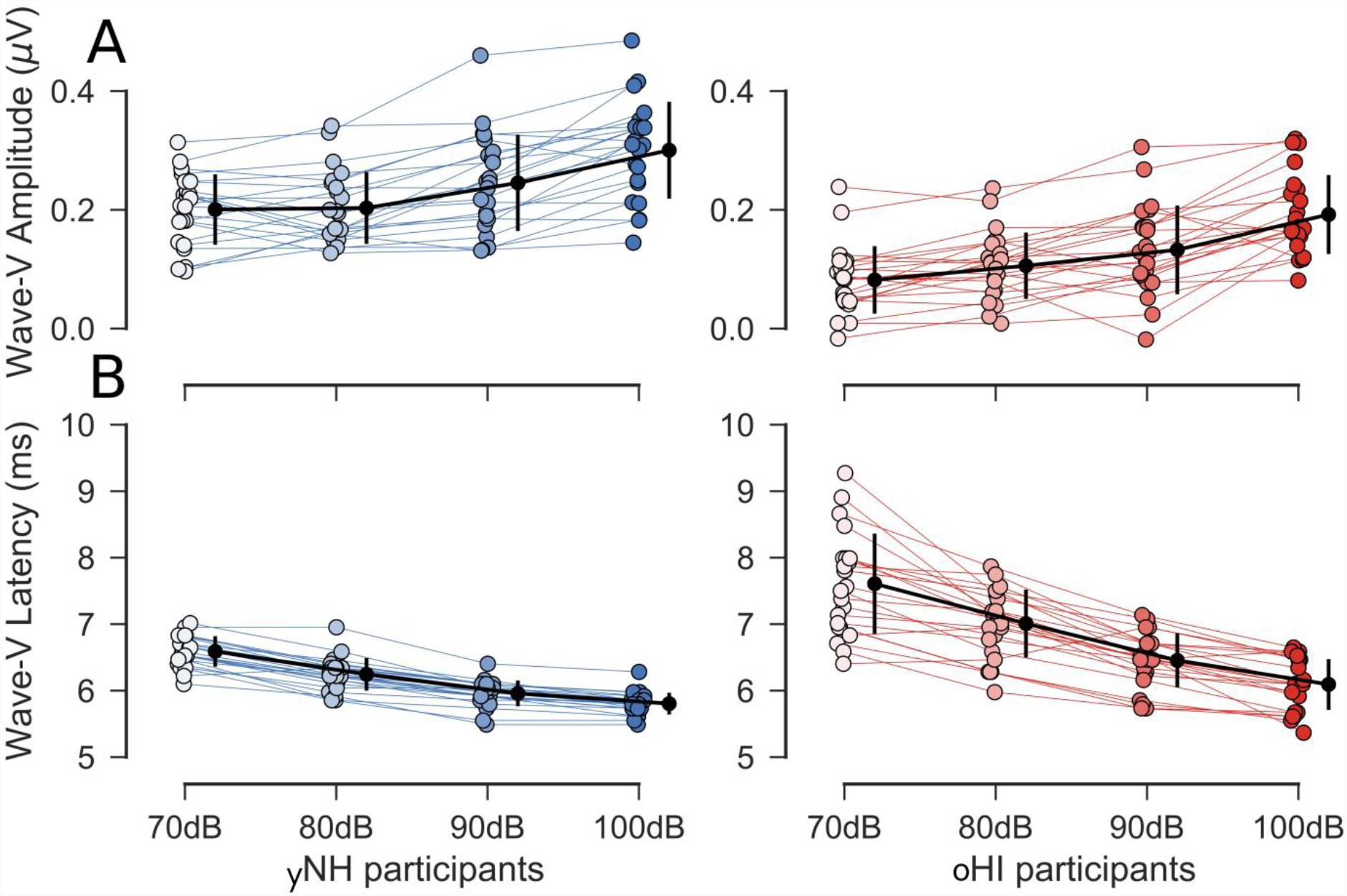
ABR Wave-V amplitude (A) and latency traces (B) for all yNH and oHI participants over the four increasing sound pressure levels (peSPLs). Error bars display mean values and standard deviation in a group per peSPL condition.

**Fig. 8:**
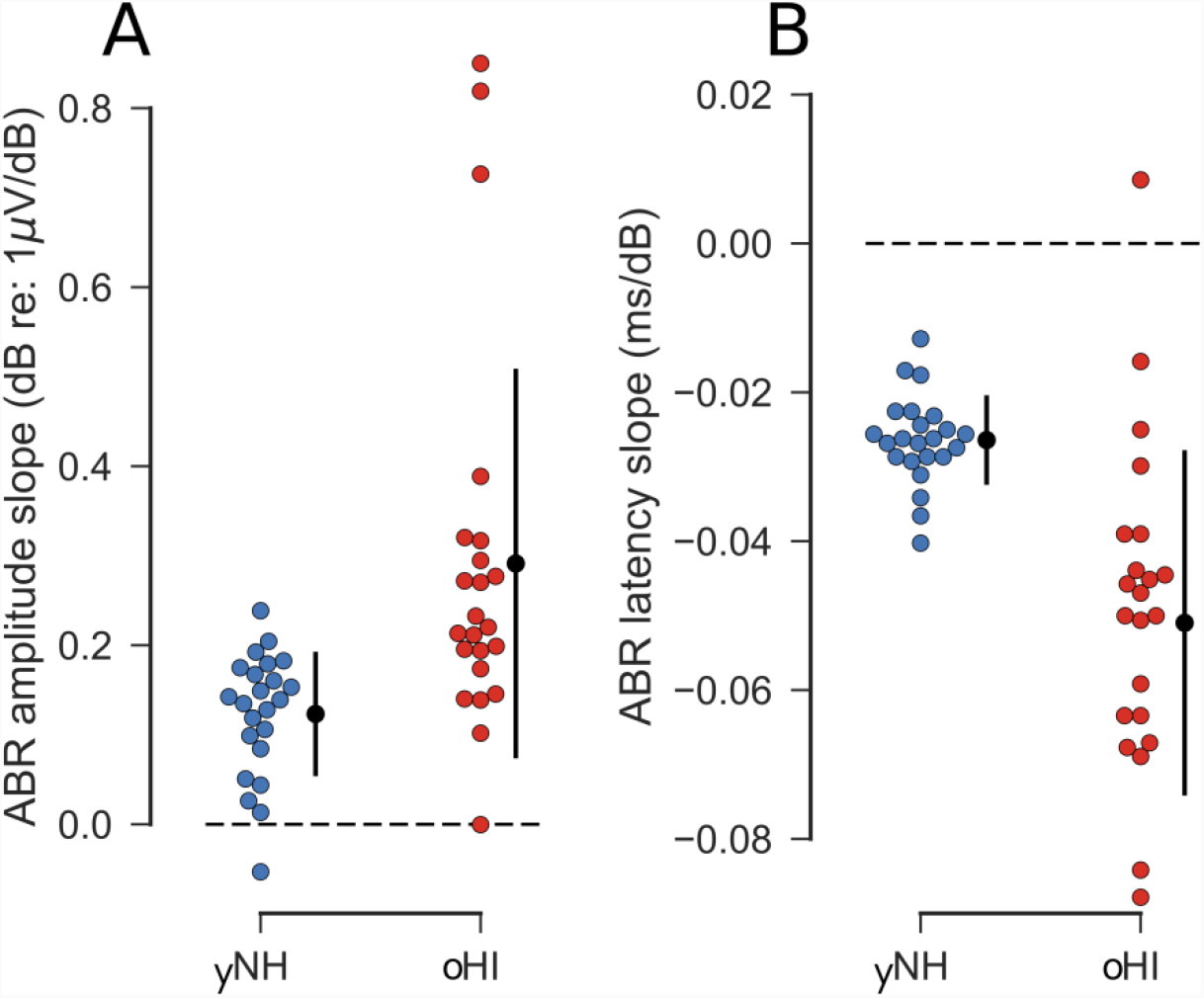
Individual ABR Wave-V (A) amplitude and (B) latency slopes (slope of straight line fitted through all four increasing peSPL conditions) for both groups (yNH, oHI). Error bars show mean and standard deviation across participants in a group. More positive amplitude slopes represent faster increasing amplitudes with increasing peSPL. More negative latency slopes represent faster decreasing latencies with increasing peSPL.

Lastly, the ratio between the ABR Wave-I and Wave-V amplitudes was computed for the 100-dB peSPL condition (see Figure 9 and 11B) as a self-normalizing measure thought to reflect central gain in the auditory brainstem (Möhrle et al., 2016; Schaette and McAlpine, 2011; Valderrama et al., 2018). The Mann-Whitney U-Test indicated no significant difference between the ABR Wave-I/V ratio of yNH and oHI participants (U = 246; p = 0.3463) suggesting the absence of a central gain mechanism in the oHI group. To investigate the gain mechanism from a different angle, we also investigated the ABR Wave-I/V ratio and its relation to the Wave-I amplitude (see Verhulst et al. (2016) for discussion). We used a multiple linear regression model predicting the Wave-I/V ratio using the Wave-I amplitude and the group as predictor variables (Figure 9). The results indicated a significant interaction term (t = −2.037, p = 0.048) aside from the expected significant effect of the Wave-I amplitude. This indicates that the slope for the yNH group is significantly shallower than that of the oHI group suggesting some degree of gain compensation in the yNH group as opposed to the oHI group (Möhrle et al., 2016).

**Fig. 9:**
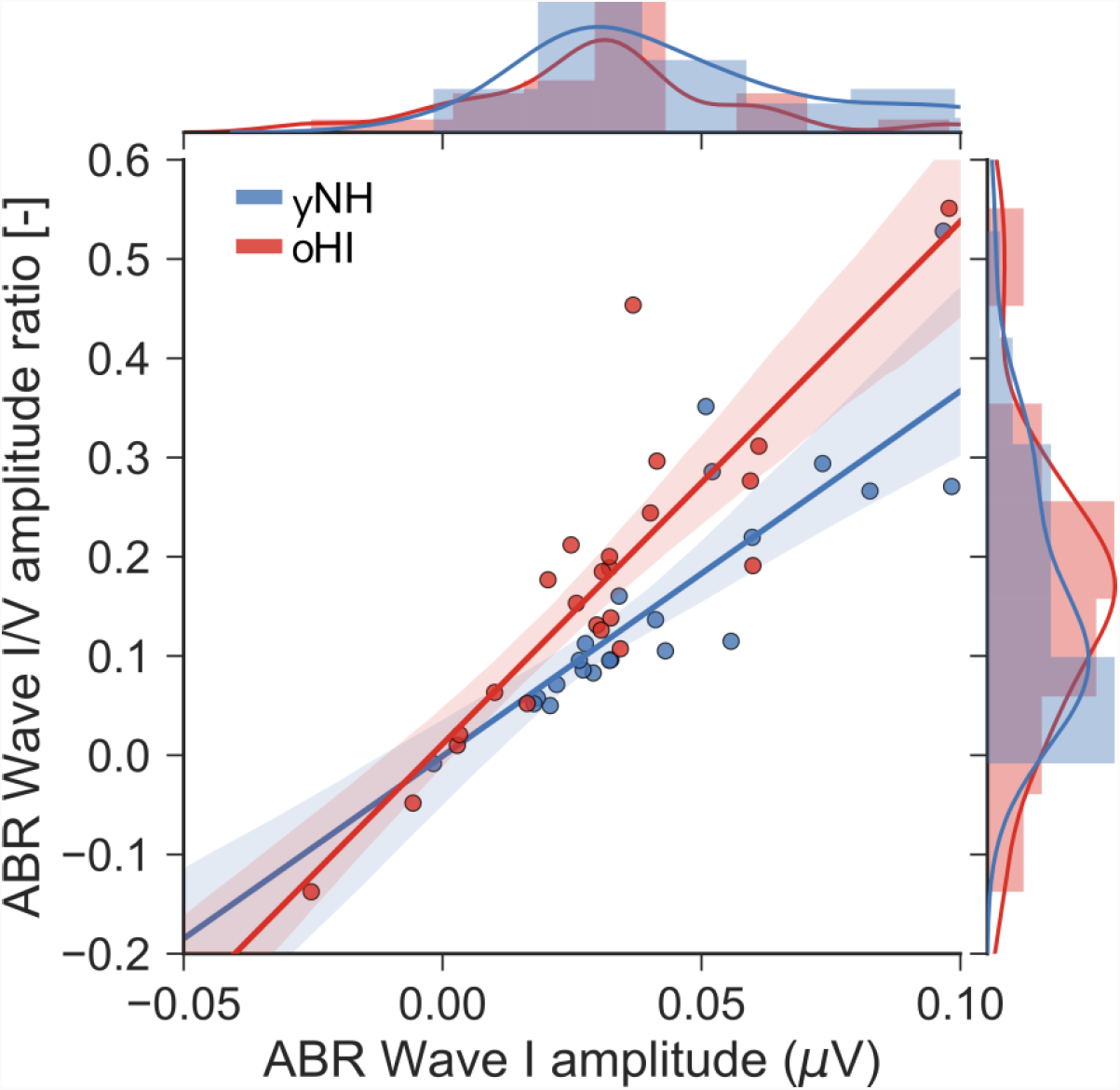
Regression plot of the ABR Wave-I amplitude with the ABR Wave-I/V amplitude slope (100 dB peSPL) for the yNH (blue) and oHI (red) participant group. The shaded areas display the 95% confidence interval of the regression fit.

### 3.4. Relationship between ABR and EFR metrics

To investigate relationships between different potential electrophysiological measures of synaptopathy, a correlation analysis was performed for the most reliable EFR (100% modulation) at 120/480 Hz and the four ABR Wave-V level conditions. The results for the 100-dB-peSPL ABR condition are visualised as regression plots in Figure 10. As expected, a significant correlation (r = 0.45; p = 0.04, N = 21) was found for the yNH participants in the BB group (f_m_:120 Hz), which elicited the strongest EFRs due to the broader excitation of the basilar membrane. A similar but weaker trend was also observed for the PT-EFR data. Even the oHI participants showed a positive trend in the BB group which was mostly driven by two participants with large EFRs. Without those two data points, the correlation disappeared completely (r = 0.0; p = 0.983, N = 17). No trends were found for the oHI-PT group. The BB-480-Hz EFRs for the yNH participants with significant responses (14 out of 21) correlated significantly with each of the four ABR level conditions (70 dB: r = 0.55, p = 0.0427; 80 dB: ρ = 0.62, p = 0.0176; 90 dB: r = 0.74, p = 0.0027; 100 dB: r = 0.58, p = 0.0301; N = 14). The relation between the BB-480-Hz EFR and the 100 dB-peSPL ABR is depicted in the first panel of Figure 10 (black data points). These correlations also held when including all data points (except for the 100-dB-peSPL ABR). If all data points (also non-significant points) were included in the BB-oHI group, a weak positive but non-significant trend was observed.

**Fig. 10:**
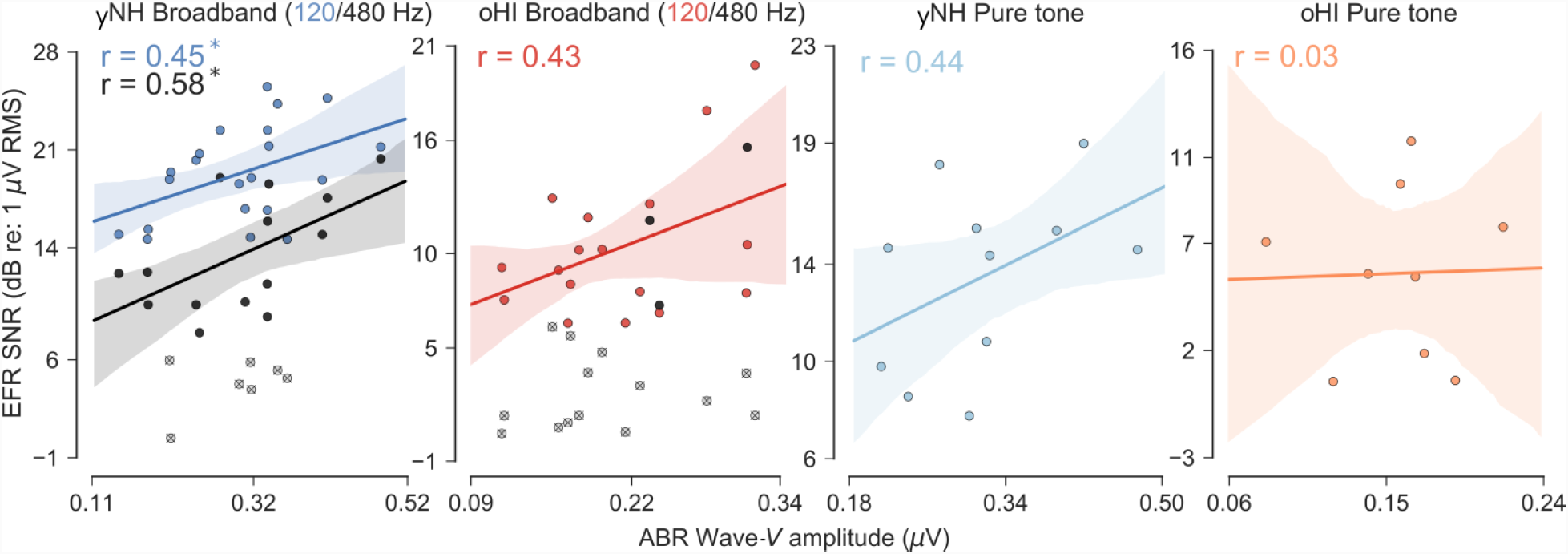
Regression plots of the ABR Wave-V amplitudes (100 dB peSPL) with the EFR SNR responses (0 dB-MD) for both stimulus conditions (BB, PT) and participant groups (yNH, oHI). For the broadband condition both available modulation frequencies 120 (colour) and 480 Hz (black) are shown. Crossed markers indicate 480-Hz EFR SNR values at noise floor level. The shaded areas display the 95% confidence interval and asterisks indicate significance (α = 0.05) of the regression fit. No regression was fitted for the oHI broadband 480-Hz condition due to too few available data points.

**Fig. 11:**
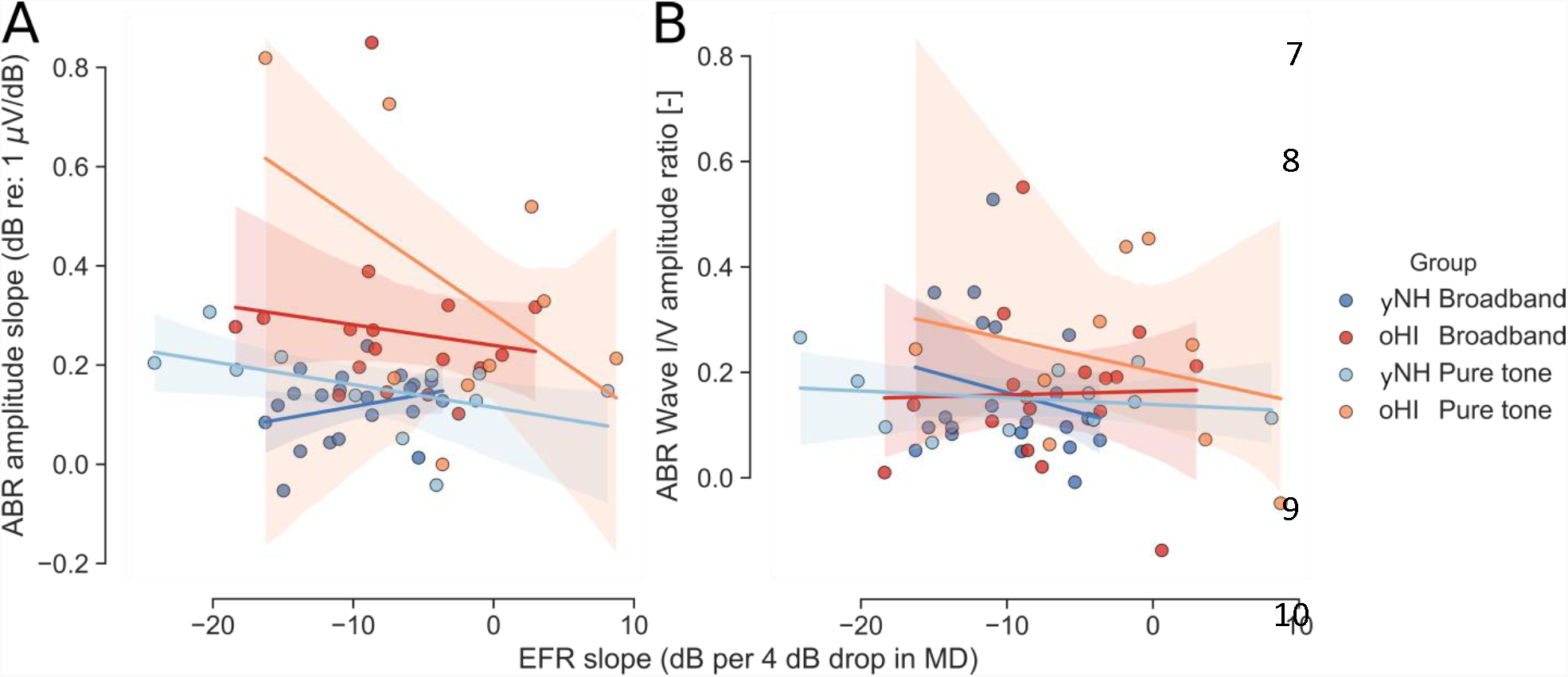
Regression plots of the EFR SNR slopes (slope of a straight line fitted through all three modulation depth conditions) with the ABR Wave-V amplitude slope (A) and the ABR Wave-I/V amplitude ratio (100 dB peSPL) for both main stimulus conditions (BB, PT) and participant groups (yNH, oHI). The shaded areas display the 95% confidence interval of the regression fit.

For the relative metrics (ABR amplitude slope and EFR slope), we expected a positive linear relationship for the yNH participants, reflecting a positive covariation of neural recruitment with temporal coding fidelity ability. However, the data did not show any significant or consistent trends (Figure 11A). As visualised in Figure 11B, the Wave-I/V ratio also did not relate to the EFR slope metric (no significance in any of the four groups). Furthermore, the 0-dB MD EFR magnitudes did not relate to the ABR amplitude slope (Figure 12). Lastly, no significant relations between the ABR Wave-I/V ratio and the 100% modulated EFR values were observed. There was only a negative trend for the BB conditions in yNH (ρ = −0.3; p = 0.1824, N = 21) and oHI (r = −0.38; p = 0.1353, N = 17) groups, who showed higher EFRs for smaller Wave-I/V ratios.

**Fig. 12:**
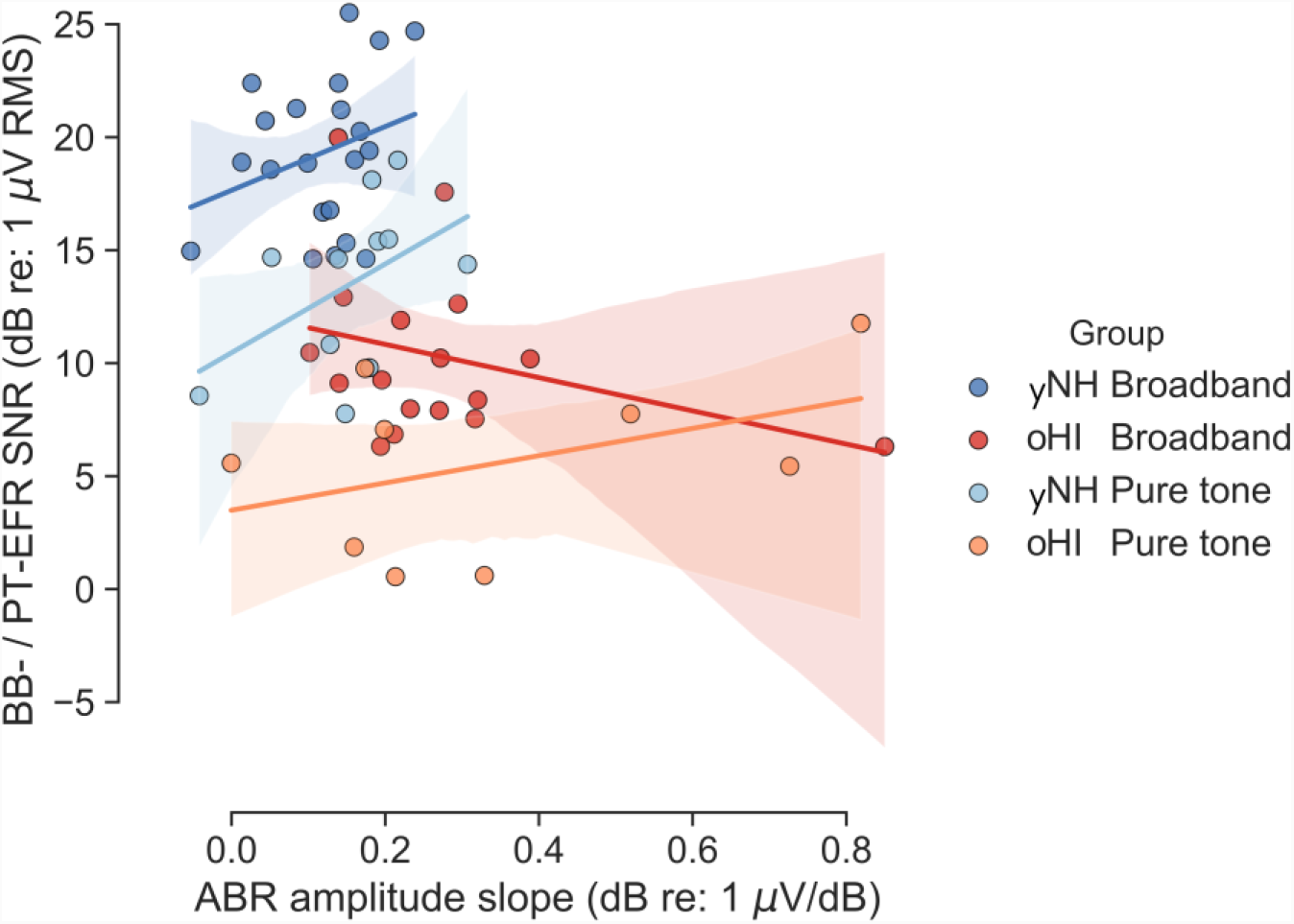
Regression plots of the ABR Wave-V amplitude slope with the 0 dB-MD EFR SNR magnitudes for both main stimulus conditions (BB, PT) and participant groups (yNH, oHI). The shaded areas display the 95% confidence interval of the regression fit.

## 4. Discussion

We measured click-ABRs at different peSPLs and EFRs to a multitude of varying stimulus parameters to contrast response behaviour of absolute and relative electrophysiological metrics in young NH and older HI participants. In general, our results show that sensorineural hearing loss has an impact on subcortical EEG measures and that their interpretation strongly depends on the underlying types of impairments at play.

### 4.1. Effects of hearing threshold elevation

#### 4.1.1. Consequences for the EFR

The auditory coding of temporal fluctuations at moderate-to-high levels is believed to be facilitated by the interplay of nerve fibres with high dynamic ranges (low-SR) and unsaturated off-frequency fibres (Encina-Llamas et al., 2017). Hearing, as tested using the classical audiogram, only requires a small amount of highly sensitive fibres (high-SR) to detect the presence of a sound (Liberman and Kujawa, 2017). A relation between audiometric thresholds and the EFR, the latter reflecting temporal coding fidelity in the early stages of the auditory system, was not expected nor found within our two participant groups (with the oHI PT-EFR condition as an exception). Nevertheless, when treating the participant sample as a whole, our data show that audiometric hearing loss does have a significant effect on EFR strength. Observing the effect only when pooling the subjects, but not within the yNH or oHI group can be explained on the basis of co-occurring synaptopathy *and* OHC deficits in oHI listeners. Hence, within a group of listeners with similar audiometric hearing loss, individual differences in EFR magnitudes would reflect individual degrees of synaptopathy. At least for the PT-EFR condition, simulations with a model of the human auditory periphery predict that individual differences in NH and HI EFRs are most impacted by different degrees of synaptopathy and to a lesser extent by OHC deficits (Verhulst et al., 2018b). EFRs were shown to reduce with decreasing modulation depth in both groups (Figure 2), implying a similar variation of the ability to encode weak temporal cues in the yNH and oHI auditory system. Our findings corroborate observations in humans (Dimitrijevic et al., 2016) and in animal models, which also show reduced EFRs with age/noise exposure (Parthasarathy et al., 2014; Parthasarathy and Kujawa, 2018; Shaheen et al., 2015).

#### 4.1.2. Relationship between SPL and the modulated neural firing rate

We contrasted subcortical EEG results of yNH and oHI participants recorded at the same SPLs to mimic every-day listening conditions. While this approach does not bring listeners with different hearing thresholds to the same sensation level (SL), it avoids the non-trivial challenge of applying an appropriate compensation method. Perceptual approaches based on the ABR/EFR stimulus threshold or on the audiogram, can introduce additional variability in the measures if the cancellation is imperfect. Additionally, it is known that the modulated firing rate of AN fibres in response to a SAM tone is not monotonously increasing with SPL but instead shows a bell shape with its maximum depending on the fibre type (low-SR or high-SR; Joris and Yin, 1992). Given a sufficient amount of intact nerve fibres, this means that a reduction of SPL within a certain range could actually lead to an increase in neural synchronisation as represented by the EFR. Applying a SL compensation might thus actually further complicate the interpretation of the data in terms of synaptopathy, instead of attempting to compensate for individual degrees of OHC deficits. If the amplification capacity of the OHCs is impaired, thereby reducing the effective drive to the IHC/AN complex, this effect of increased modulated firing rates for reduced SPL input, might also be observable in EFRs to medium-to-high stimulus SPLs. This mechanism might partly explain the significantly increasing yNH-EFR magnitudes in response to the 5 dB stimulus level decrease for otherwise identical narrowband conditions (Figure 5A). The same mechanism could also have been the basis of the significant positive correlation between oHI PT-EFRs and the audiogram threshold at 4 kHz. In line with our findings, enhanced sensitivity to amplitude modulation, after the introduction of noise-induced hearing loss, was also reported for chinchillas for carrier frequencies above 2 kHz and similar modulation frequencies (Zhong et al., 2014). The authors of the latter study interpreted this as a compensatory mechanism of the periphery and/or central structures.

#### 4.1.3. Consequences for the ABR

While showing good retest-reliability at medium SPL in NH participants (Prendergast et al., 2018), the Wave I in humans is harder to measure than the Wave V using the vertex configuration and varies greatly in amplitude and latency between participants (Beattie, 1988; Lauter and Loomis, 1988; Mehraei et al., 2016; Trune et al., 1988). Especially in the oHI group, the ABR Wave I at lower SPLs were often too weak to be reliably extracted from the recordings. It can therefore only be suspected that the ABR Wave-I amplitude for the oHI group was strongly reduced by high-frequency OHC loss as computational modelling work has predicted (Verhulst et al., 2016). On the other hand, the ABR Wave V is very robust in humans and is believed to be generated in the lateral lemniscus and inferior colliculus (Melcher et al., 1996; Møller and Jannetta, 1985). The ABR Wave V was clearly detectable from our recordings even for oHI participants (Figure 6/7). In agreement with earlier observations (Konrad-Martin et al., 2012) elevated hearing thresholds resulted in reduced ABR Wave-V amplitudes in all tested SPL conditions in comparison to the yNH-ABRs.

Even though the Wave V is a marker of very early auditory processing, it cannot deliver identical information to the Wave I regarding AN processing. It is therefore important to consider whether the use of Wave V can be discussed in the context of synaptopathy. At least several aspects of Wave-I characteristics are reflected in Wave V. For example, changes in ABR Wave-I amplitude have been shown to be mirrored in the Wave-V latency shift for varying masking noise levels while ABR Wave-I and V amplitudes did not relate in the quiet condition without masking noise (Mehraei et al., 2016). It was also reported that the latencies of both waves covary for frequency contributions above 2 kHz (Don and Eggermont, 1978). Based on this observation, and the particularly strong vulnerability of high-frequency cochlear regions to OHC loss, the increased Wave-V latencies in oHI participants might be interpreted as a neural marker for high-frequency OHC loss.

In contrast to the earlier waves, Don & Eggermont (1978) also showed that the ABR Wave-V amplitude is independent from the frequency regions contributing to its generation. It was also reported that the Wave-V and Wave-IV amplitudes are not directly related to the amount of cochlear synaptopathy in mice (Möhrle et al., 2016; Sergeyenko et al., 2013). These findings suggest that the reduced Wave-V peak amplitudes we reported for the oHI group might predominantly be attributed to OHC loss. Nevertheless, a reduction in the number of nerve fibres as a consequence of cochlear synaptopathy cannot be excluded as a possible contributor as functional ABR model simulations show that both, synaptopathy and OHC loss, might reduce the Wave-V amplitude (Verhulst et al., 2016) in the absence of compensatory brainstem gain mechanisms (Chambers et al., 2016; Möhrle et al., 2016).

### 4.2. Effects of stimulus features

#### 4.2.1. Effects of bandwidth on the EFR

Our results show clear differences as a consequence of manipulations of the stimulus bandwidth in both groups (Figure 5A). An expected significant decrease in EFR magnitude with stimulus bandwidth reduction was only observed in the yNH group and might reflect a more restricted synchronised firing of the spiral ganglion cells and subsequent neural processing stages in the ascending auditory pathway. Unfortunately, the PT conditions did not result in maximising individual differences, but rather led to a greater number of EFRs at noise floor level in the oHI participants. The NB stimulus (at least at 75 dB SPL) was able to evoke significant responses in both groups while retaining good frequency specificity. However, without the use of off-frequency masking, the relative contributions of off-frequency fibres to the NB-EFR response remains unclear. The lack of sensitivity to the stimulus-bandwidth changes in the oHI-EFRs might be attributed to the wider auditory filters as a consequence of OHC loss. A wider band of frequencies falls within a certain filter, thereby reducing the frequency selectivity and associated frequency-specific coding. In addition to the consequences of OHC loss, a reduction of the number of nerve fibres might lead to a diminished ability of the brain to code subtle stimulus envelope changes robustly enough to be picked up by the scalp EEG.

#### 4.2.2. Effects of modulation frequency and the EFR generators

Increasing the modulation frequency of the stimulus reduced the EFR significantly and similarly for both groups (Figure 5B). Our results corroborate the EFR reductions observed for higher modulation frequencies in temporal modulation transfer functions (Purcell et al., 2004). Because we compensated for the frequency dependent noise floor (~1/f) by using the EFR SNR metric, the effect of modulation frequency on the EFR is likely attributed to the frequency dependent constructive and destructive phase interferences of multiple neural generators thought to be responsible for the EFR generation (Tichko and Skoe, 2017). Even though it is much harder to measure the EFR to a 480-Hz modulator, they might reflect more peripheral generators than the IC and offer a better proxy measure for synaptopathy (Shaheen et al., 2015). Our finding that the BB-EFR at 480 Hz for yNH participants is more strongly related to the ABR Wave-V measure compared to the BB-EFR at 120 Hz (Figure 10) is in line with this idea. But, the higher modulation frequency comes at the cost of missing data points due to generally weak responses in oHI group. Note that the argument about more peripheral sources for higher modulation frequencies (Purcell et al., 2004) would only hold for noise carriers. The use of a 480-Hz modulator on a pure tone carrier yields resolved AM side-bands (Kohlrausch et al., 2000), which result in an across-frequency spectral representation of the AM stimulus rather than a temporal envelope cue associated with AM side-band frequencies that fall within a single auditory filter.

### 4.3. Applicability of relative metrics in the normal and impaired auditory system

#### 4.3.1. The EFR slope

Our yNH BB-EFR slope metric (Figure 3) showed comparable values to those reported in a previous study (for a 100 Hz transposed 4 kHz pure tone carrier; Bharadwaj et al., 2015). The latter study related shallower EFR slopes to better AM and ITD detection thresholds as well as selective attention task performance. Different from that study, we did not use off-frequency notched noise masking. Consequently, the wider spread in the slope values in the PT compared to the BB condition could in part be explained by an additional off-frequency contribution to the PT-EFRs, whose extent might vary between listeners. We did not observe significant differences in the steepness of the EFR slopes between the yNH and oHI group in contrast to what we hypothesised. This challenges the interpretation of the EFR slope measure in light of synaptopathy, as it is expected that synaptopathy occurs before OHC loss sets in with age (Parthasarathy and Kujawa, 2018; Sergeyenko et al., 2013). However, the potential contribution of off-frequency fibres to the PT-EFRs might have affected the slope metrics in the two groups differently and thereby washed out the expected effects caused by synaptopathy; i.e. the increased cochlear filter width in the oHI population could have yielded stronger off-frequency contributions compared to the yNH group. However, the same mechanism could not have been responsible for the missing EFR slope differences in the BB conditions between groups due to the broadband characteristic of the stimulus. We therefore conclude that factors such as OHC loss and synaptopathy have different impacts on the EFR slope metric and that it is not exclusively sensitive to synaptopathy. The potential presence of neural *and* sensory factors of peripheral hearing loss on the EFR slope metric does not allow for a clear interpretation of this metric in the oHI group. The few (mainly oHI participants) showing positive slope values reflect the susceptibility of the EFR slope metric to outliers and high variability and could be partially explained by potential non-linearities in the AM depth function (Dimitrijevic et al., 2016).

#### 4.3.2. The ABR amplitude and latency slope

Our data revealed shallower ABR amplitude and latency slopes for the yNH group in contrast to the oHI group for peSPL levels between 70 and 100 dB (Figure 8). At those stimulus levels, low-SR fibres will already have reached their maximum discharge rate and any further increase in amplitude of the brainstem response might only be due to the contributions of low-SR fibres that show higher saturation levels (Heinz and Young, 2004; Liberman, 1978). A steep slope in the yNH group might therefore reflect a healthy SR-fibre population consisting of both low and high SR fibres (Furman et al., 2013). However, in contrast to our expectations based on the hypothesis that synaptopathy is the main driving force of this metric, the oHI group actually showed steeper ABR Wave-V amplitude growth in agreement with Gorga et al. (1985). OHC loss therefore seems to play the dominating role in determining the amplitude slope, by showing a recruitment of additional high-frequency channels with increasing peSPLs which result in stronger synchronised responses and steeper amplitude slopes. This broadening of peripheral auditory filters at high SPLs results in a more basally peaking excitation pattern (Ren, 2002) which explains the steeper oHI-ABR latency slopes we found for listeners with steep ABR amplitude slopes. Our findings of increased latencies and steeper ABR latency slopes in the oHI participants are in line with other studies (e.g. Burkard and Sims, 2002; Prendergast et al., 2017) who described an increase of Wave-V latency with age or lifetime noise exposure and as a function of audiometrically sloping hearing loss. The yNH latency slopes were less steep than those reported for the oHI group and corroborate slope values reported in many other studies (e.g. Dau, 2003; Don and Eggermont, 1978; Eggermont and Don, 1980; Mehraei et al., 2016). At 100 dB SPL, the latencies of both groups converged and no longer differed significantly (Figure 7). At this high stimulus level, latency and amplitude actually show a significant negative relationship in the oHI group indicating that at high peSPL, the ABR Wave-V amplitude is proportional to the extent of the broadening auditory filters caused by OHC loss. These results suggest that increased stimulus levels can restore the oHI-ABR latency to normal while the amplitudes remain smaller in this group (Lewis et al., 2015; Neely et al., 2003).

The impossibility of restoring the ABR amplitude at higher stimulus levels might be indicative of additional neural fibre loss in the oHI group. Participants that have low latency values at high peSPLs (normal OHC function) whilst also showing low amplitude values (low synchronised neural response) might be impacted particularly strongly by cochlear synaptopathy (Verhulst et al., 2016). This idea stands in contrast to the hypothesis of a central gain compensation mechanism (Chambers et al., 2016; Schaette and McAlpine, 2011). Our data did not show differences between the yNH and oHI group for the Wave-I/V ratio as a potential marker of such a neural gain mechanism at the level of the brainstem. Given the degraded auditory input for the oHI participants, a smaller Wave-I/V ratio would have been expected (Schaette and McAlpine, 2011). Nevertheless, the shallower slopes in the yNH group contrasting the Wave-I amplitude against the Wave-I/V ratio (Figure 9) might be indicative of some degree of gain compensation in the yNH group (see Verhulst et al., 2016 for a discussion of underlying effects). The missing gain effect in the oHI group might be partially explained by considering age as a factor. It was recently shown that such a compensatory gain mechanism was present only in aged, not old, rats in a similar SPL range as was considered here (Möhrle et al., 2016).

### 4.4. Interrelation of EFR and ABR metrics

Contrasting the normal and impaired auditory system, we found a significant relation between the yNH-ABR Wave V and yNH-EFR amplitudes for the 120 and 480 Hz modulated BB condition (Figure 10). As the neurons responsible for coding temporal stimulus modulations are presumably only a subset of all neurons responding to the ABR click stimulus, this relationship between overlapping neural generators was expected. Interestingly, this relationship breaks down in participants with peripheral hearing loss. Other authors who reported similar relationships between the EFR and ABR in rats interpret this as a decoupling of phasic and tonic synchrony caused by changing ratios of neurons representing the different stimuli with age (Parthasarathy et al., 2014). Another potential mechanism that could have caused the missing relation between the oHI-ABR and oHI-EFR is the loss of frequency selectivity that would allow more neurons to respond to the AM stimuli, without altering the neural contributions to the broadband click ABR.

The missing correlations between the EFR slope as a normalised measure of temporal coding fidelity and the ABR amplitude slope as a normalised metric of neural recruitment in both groups was unexpected, as other studies have suggested that both metrics independently relate to synaptopathy expression (e.g. Bharadwaj et al., 2014; Furman et al., 2013). Similarly, if central gain and cochlear synaptopathy are two simultaneously occurring phenomena in the peripheral auditory system, we should see a correlation between the ABR Wave-I/V ratio and the EFR slope metric. This correlation should be particularly pronounced for the oHI group given their degraded auditory input due to OHC and neural fibre loss. This was not the case in our data (Figure 11B).

The missing relationships could have several reasons. The ABR amplitude slopes in the yNH group showed very homogenous values and therefore very little variance. The degree to which the participants in this study are affected by nerve fibre loss might just be too small to be uncovered by the presented metrics. A similar reasoning could be applied to the yNH Wave-I/V ratio. Despite findings in other studies (Schaette and McAlpine, 2011; Valderrama et al., 2018), the Wave-I/V ratio might not be a very reliable measure of central gain, given the difficult ascertainment of the Wave I in humans, especially for HI participants. In this context, other groups have also failed to show a difference between normal-hearing and tinnitus participants with normal audiometric thresholds (Guest et al., 2017). As described earlier, the EFR slope is hard to interpret in the presence of sensorineural hearing loss. Similarly, the oHI-ABR amplitude slope might primarily reflect OHC loss which is not directly linked to cochlear synaptopathy.

### 4.5. Suitability for clinical applications

In contrast to other groups (e.g. Bramhall et al., 2017; Liberman et al., 2016) who recorded Wave I with gold foil tiptrode electrodes in NH groups, our ABR Wave-I amplitudes were recorded with a conventional EEG setup and were often too weak to be picked up reliably. For example, Wave-I data for our oHI group did not show any consistent trajectory across peSPL conditions (Figure 6). Due to its difficult ascertainment and high variability using standard procedures, this measure is less attractive for diagnostic purposes in the context of synaptopathy and has already been shown to be an unreliable predictor for hearing-in-noise performance (Lobarinas et al., 2017). A big caveat of the use of EFRs in a clinical context is the low signal strength, primarily for the oHI group. Particularly, for the pure tone conditions, lower modulation depths and high modulation frequencies yielded EFRs that could not be distinguished from the noise floor. Even though the pure tone and narrowband stimuli deliver more frequency specific information about potential fibre loss as compared to the broadband stimuli, a quantitative interpretation of EFR magnitudes in relation to synaptopathy is problematic without an appropriate reference condition. Future studies should improve the EFR stimulus design to achieve stronger and more frequency specific EFRs.

## 5. Summary and Conclusion

The present study investigated differences in multiple subcortical EEG measures in young NH and older HI participant groups to investigate how these metric behave in the presence of sensory and neural hearing deficits. We showed that EFRs and ABRs can be recorded in aged participants with a high degree of sloping sensorineural hearing loss for certain stimulus conditions. Both ABRs and EFRs showed lower amplitudes in the oHI group and were only significantly related in the yNH group, thereby replicating the findings from numerous other human and animal studies. The use of relative ABR and EFR metrics can provide valuable insight into the underlying hearing deficit mechanisms but the high degree of variability (especially in the oHI group) allows interpretations only on a group level and is, in the current design, less suitable for individual diagnostics. The ambiguity in interpreting the oHI findings in the presence of peripheral sensory *and* neural hearing loss underlines the suggestions of other authors (e.g. Kobel et al., 2017; Plack et al., 2016) who argue that a reliable diagnostic procedure for disentangling different contributors must involve a battery of electrophysiological *and* behavioural tests to reduce the likelihood of incorrect diagnoses. This study reported what, realistically, can be expected when applying and combining different established electrophysiological metrics to the normal and aged impaired auditory system. Our findings can help guide future developments of electrophysiological measures of cochlear synaptopathy toward becoming a reliable and informative clinical tool.

## Acknowledgement

The authors thank Anoop Jagadeesh, Frauke Ernst, Marvin Schmidt and Moritz Wächtler for help with the data collection.

## References

Beattie, R.C., 1988. Interaction of click polarity, stimulus level, and repetition rate on the auditory brainstem response. Scand Audiol 17, 99–109. https://doi.org/10.3109/01050398809070698

Bharadwaj, H.M., Masud, S., Mehraei, G., Verhulst, S., Shinn-Cunningham, B.G., 2015. Individual differences reveal correlates of hidden hearing deficits. J. Neurosci. 35, 2161–2172. https://doi.org/10.1523/JNEUROSCI.3915-14.2015

Bharadwaj, H.M., Verhulst, S., Shaheen, L., Liberman, M.C., Shinn-Cunningham, B.G., 2014. Cochlear neuropathy and the coding of supra-threshold sound. Front Syst Neurosci 8, 26. https://doi.org/10.3389/fnsys.2014.00026

Bourien, J., Tang, Y., Batrel, C., Huet, A., Lenoir, M., Ladrech, S., Desmadryl, G., Nouvian, R., Puel, J.-L., Wang, J., 2014. Contribution of auditory nerve fibers to compound action potential of the auditory nerve. J. Neurophysiol. 112, 1025–1039. https://doi.org/10.1152/jn.00738.2013

Bramhall, N.F., Konrad-Martin, D., McMillan, G.P., Griest, S.E., 2017. Auditory Brainstem Response Altered in Humans With Noise Exposure Despite Normal Outer Hair Cell Function. Ear Hear 38, e1–e12. https://doi.org/10.1097/AUD.0000000000000370

Burkard, R.F., Sims, D., 2002. A comparison of the effects of broadband masking noise on the auditory brainstem response in young and older adults. Am J Audiol 11, 13–22. https://doi.org/10.1044/1059-0889(2002/004)

Carney, L.H., 2018. Supra-Threshold Hearing and Fluctuation Profiles: Implications for Sensorineural and Hidden Hearing Loss. J. Assoc. Res. Otolaryngol. https://doi.org/10.1007/s10162-018-0669-5

Chambers, A.R., Resnik, J., Yuan, Y., Whitton, J.P., Edge, A.S., Liberman, M.C., Polley, D.B., 2016. Central Gain Restores Auditory Processing following Near-Complete Cochlear Denervation. Neuron 89, 867–879. https://doi.org/10.1016/j.neuron.2015.12.041

Chen, G.-D., Tanaka, C., Henderson, D., 2008. Relation between outer hair cell loss and hearing loss in rats exposed to styrene. Hear. Res. 243, 28–34. https://doi.org/10.1016/j.heares.2008.05.008

Dau, T., 2003. The importance of cochlear processing for the formation of auditory brainstem and frequency following responses. J. Acoust. Soc. Am. 113, 936–950. https://doi.org/10.1121/1.1534833

Debener, S., Thorne, J., Schneider, T.R., Viola, F.C., 2010. Using ICA for the Analysis of Multi-Channel EEG Data, in: Ullsperger, M., Debener, S. (Eds.), Simultaneous EEG and FMRI: Recording, Analysis, and Application. Oxford University Press, pp. 121–133.

Dimitrijevic, A., Alsamri, J., John, M.S., Purcell, D., George, S., Zeng, F.-G., 2016. Human Envelope Following Responses to Amplitude Modulation: Effects of Aging and Modulation Depth. Ear Hear 37, e322–335. https://doi.org/10.1097/AUD.0000000000000324

Don, M., Eggermont, J.J., 1978. Analysis of the click-evoked brainstem potentials in man using high-pass noise masking. J. Acoust. Soc. Am. 63, 1084–1092. https://doi.org/10.1121/1.381816

Eggermont, J.J., Don, M., 1980. Analysis of the click-evoked brainstem potentials in humans using high-pass noise masking. II. Effect of click intensity. J. Acoust. Soc. Am. 68, 1671–1675. https://doi.org/10.1121/1.385199

Encina-Llamas, G., Parthasarathy, A., Harte, J.M., Dau, T., Kujawa, S.G., Shinn-Cunningham, B., Epp, B., 2017. Hidden hearing loss with envelope following responses (EFRs): The off-frequency problem.

Fernandez, K.A., Jeffers, P.W.C., Lall, K., Liberman, M.C., Kujawa, S.G., 2015. Aging after noise exposure: acceleration of cochlear synaptopathy in “recovered” ears. J. Neurosci. 35, 7509–7520. https://doi.org/10.1523/JNEUROSCI.5138-14.2015

Festen, J.M., Plomp, R., 1983. Relations between auditory functions in impaired hearing. J. Acoust. Soc. Am. 73, 652–662. https://doi.org/10.1121/1.388957

Fulbright, A.N.C., Le Prell, C.G., Griffiths, S.K., Lobarinas, E., 2017. Effects of Recreational Noise on Threshold and Suprathreshold Measures of Auditory Function. Semin Hear 38, 298–318. https://doi.org/10.1055/s-0037-1606325

Furman, A.C., Kujawa, S.G., Liberman, M.C., 2013. Noise-induced cochlear neuropathy is selective for fibers with low spontaneous rates. J. Neurophysiol. 110, 577–586. https://doi.org/10.1152/jn.00164.2013

Glasberg, B.R., Moore, B.C., 1986. Auditory filter shapes in subjects with unilateral and bilateral cochlear impairments. J. Acoust. Soc. Am. 79, 1020–1033. https://doi.org/10.1121/1.393374

Gorga, M.P., Worthington, D.W., Reiland, J.K., Beauchaine, K.A., Goldgar, D.E., 1985. Some comparisons between auditory brain stem response thresholds, latencies, and the pure-tone audiogram. Ear Hear 6, 105–112. https://doi.org/10.1097/00003446-198503000-00008

Gramfort, A., Luessi, M., Larson, E., Engemann, D.A., Strohmeier, D., Brodbeck, C., Goj, R., Jas, M., Brooks, T., Parkkonen, L., Hämäläinen, M., 2013. MEG and EEG data analysis with MNE-Python. Front Neurosci 7, 267. https://doi.org/10.3389/fnins.2013.00267

Gramfort, A., Luessi, M., Larson, E., Engemann, D.A., Strohmeier, D., Brodbeck, C., Parkkonen, L., Hämäläinen, M.S., 2014. MNE software for processing MEG and EEG data. Neuroimage 86, 446–460. https://doi.org/10.1016/j.neuroimage.2013.10.027

Guest, H., Munro, K.J., Prendergast, G., Howe, S., Plack, C.J., 2017. Tinnitus with a normal audiogram: Relation to noise exposure but no evidence for cochlear synaptopathy. Hear. Res. 344, 265–274. https://doi.org/10.1016/j.heares.2016.12.002

Heinz, M.G., Young, E.D., 2004. Response growth with sound level in auditory-nerve fibers after noise-induced hearing loss. J. Neurophysiol. 91, 784–795. https://doi.org/10.1152/jn.00776.2003

Hickox, A.E., Larsen, E., Heinz, M.G., Shinobu, L., Whitton, J.P., 2017. Translational issues in cochlear synaptopathy. Hear. Res. 349, 164–171. https://doi.org/10.1016/j.heares.2016.12.010

Hind, S.E., Haines-Bazrafshan, R., Benton, C.L., Brassington, W., Towle, B., Moore, D.R., 2011. Prevalence of clinical REFerrals having hearing thresholds within normal limits. Int J Audiol 50, 708–716. https://doi.org/10.3109/14992027.2011.582049

Humes, L.E., 2013. Understanding the speech-understanding problems of older adults. Am J Audiol 22, 303–305. https://doi.org/10.1044/1059-0889(2013/12-0066)

Humes, L.E., Kewley-Port, D., Fogerty, D., Kinney, D., 2010. Measures of Hearing Threshold and Temporal Processing across the Adult Lifespan. Hear Res 264, 30–40. https://doi.org/10.1016/j.heares.2009.09.010

ISO 1990. Acoustics - Determination of occupational noise exposure and estimation of noise-induced hearing impairment. International Organization of Standardization, ISO 1999:1990, Switzerland.

Johnson, E.W., 1970. Tuning forks to audiometers and back again. Laryngoscope 80, 49–68. https://doi.org/10.1288/00005537-197001000-00005

Joris, P.X., Yin, T.C., 1992. Responses to amplitude-modulated tones in the auditory nerve of the cat. J. Acoust. Soc. Am. 91, 215–232. https://doi.org/10.1121/1.402757

Kobel, M., Le Prell, C.G., Liu, J., Hawks, J.W., Bao, J., 2017. Noise-induced cochlear synaptopathy: Past findings and future studies. Hear. Res. 349, 148–154. https://doi.org/10.1016/j.heares.2016.12.008

Kohlrausch, A., Fassel, R., Dau, T., 2000. The influence of carrier level and frequency on modulation and beat-detection thresholds for sinusoidal carriers. J. Acoust. Soc. Am. 108, 723–734.

Konrad-Martin, D., Dille, M.F., McMillan, G., Griest, S., McDermott, D., Fausti, S.A., Austin, D.F., 2012. Age-related changes in the auditory brainstem response. J Am Acad Audiol 23, 18–75. https://doi.org/10.3766/jaaa.23.1.3

Kujawa, S.G., Liberman, M.C., 2009. Adding insult to injury: cochlear nerve degeneration after “temporary” noise-induced hearing loss. J. Neurosci. 29, 14077–14085. https://doi.org/10.1523/JNEUROSCI.2845-09.2009

Kumar, G., Amen, F., Roy, D., 2007. Normal hearing tests: is a further appointment really necessary? J R Soc Med 100, 66. https://doi.org/10.1177/014107680710000212

Lauter, J.L., Loomis, R.L., 1988. Individual differences in auditory electric responses: comparisons of between-subject and within-subject variability. II. Amplitude of brainstem Vertex-positive peaks. Scand Audiol 17, 87–92. https://doi.org/10.3109/01050398809070696

Lewis, J.D., Kopun, J., Neely, S.T., Schmid, K.K., Gorga, M.P., 2015. Tone-burst auditory brainstem response wave V latencies in normal-hearing and hearing-impaired ears. J. Acoust. Soc. Am. 138, 3210–3219. https://doi.org/10.1121/1.4935516

Liberman, M.C., 1978. Auditory-nerve response from cats raised in a low-noise chamber. J. Acoust. Soc. Am. 63, 442–455. https://doi.org/10.1121/1.381736

Liberman, M.C., Epstein, M.J., Cleveland, S.S., Wang, H., Maison, S.F., 2016. Toward a Differential Diagnosis of Hidden Hearing Loss in Humans. PLoS ONE 11, e0162726. https://doi.org/10.1371/journal.pone.0162726

Liberman, M.C., Kujawa, S.G., 2017. Cochlear synaptopathy in acquired sensorineural hearing loss: Manifestations and mechanisms. Hear. Res. 349, 138–147. https://doi.org/10.1016/j.heares.2017.01.003

Lin, H.W., Furman, A.C., Kujawa, S.G., Liberman, M.C., 2011. Primary Neural Degeneration in the Guinea Pig Cochlea After Reversible Noise-Induced Threshold Shift. J Assoc Res Otolaryngol 12, 605–616. https://doi.org/10.1007/s10162-011-0277-0

Lobarinas, E., Spankovich, C., Le Prell, C.G., 2017. Evidence of “hidden hearing loss” following noise exposures that produce robust TTS and ABR wave-I amplitude reductions. Hear. Res. 349, 155–163. https://doi.org/10.1016/j.heares.2016.12.009

Makary, C.A., Shin, J., Kujawa, S.G., Liberman, M.C., Merchant, S.N., 2011. Age-related primary cochlear neuronal degeneration in human temporal bones. J. Assoc. Res. Otolaryngol. 12, 711–717. https://doi.org/10.1007/s10162-011-0283-2

Mehraei, G., Hickox, A.E., Bharadwaj, H.M., Goldberg, H., Verhulst, S., Liberman, M.C., Shinn-Cunningham, B.G., 2016. Auditory Brainstem Response Latency in Noise as a Marker of Cochlear Synaptopathy. J. Neurosci. 36, 3755–3764. https://doi.org/10.1523/JNEUROSCI.4460-15.2016

Melcher, J.R., Knudson, I.M., Fullerton, B.C., Guinan, J.J., Norris, B.E., Kiang, N.Y., 1996. Generators of the brainstem auditory evoked potential in cat. I. An experimental approach to their identification. Hear. Res. 93, 1–27. https://doi.org/10.1016/0378-5955(95)00178-6

Millman, K.J., Aivazis, M., 2011. Python for Scientists and Engineers. Computing in Science Engineering 13, 9–12. https://doi.org/10.1109/MCSE.2011.36

Mitchell, C., Phillips, D.S., Trune, D.R., 1989. Variables affecting the auditory brainstem response: audiogram, age, gender and head size. Hear. Res. 40, 75–85. https://doi.org/10.1016/0378-5955(89)90101-9

Möhrle, D., Ni, K., Varakina, K., Bing, D., Lee, S.C., Zimmermann, U., Knipper, M., Rüttiger, L., 2016. Loss of auditory sensitivity from inner hair cell synaptopathy can be centrally compensated in the young but not old brain. Neurobiol. Aging 44, 173–184. https://doi.org/10.1016/j.neurobiolaging.2016.05.001

Møller, A., Jannetta, P.J., 1985. Neural Generators of the Auditory Brainstem Response, in: Jacobson, J.T. (Ed.), The Auditory Brainstem Response. College Hill, San Diego, pp. 13–31.

Neely, S.T., Gorga, M.P., Dorn, P.A., 2003. Cochlear compression estimates from measurements of distortion-product otoacoustic emissions. J. Acoust. Soc. Am. 114, 1499–1507. https://doi.org/10.1121/1.1604122

Oliphant, T.E., 2007. Python for Scientific Computing. Computing in Science Engineering 9, 10–20. https://doi.org/10.1109/MCSE.2007.58

Parthasarathy, A., Datta, J., Torres, J.A.L., Hopkins, C., Bartlett, E.L., 2014. Age-related changes in the relationship between auditory brainstem responses and envelope-following responses. J. Assoc. Res. Otolaryngol. 15, 649–661. https://doi.org/10.1007/s10162-014-0460-1

Parthasarathy, A., Kujawa, S.G., 2018. Synaptopathy in the Aging Cochlea: Characterizing Early-Neural Deficits in Auditory Temporal Envelope Processing. J. Neurosci. 38, 7108–7119. https://doi.org/10.1523/JNEUROSCI.3240-17.2018

Pinheiro, J., Bates, D., DebRoy, S., Sarkar, D., R Core Team, 2017. nlme: Linear and Nonlinear Mixed Effects Models.

Plack, C.J., Léger, A., Prendergast, G., Kluk, K., Guest, H., Munro, K.J., 2016. Toward a Diagnostic Test for Hidden Hearing Loss. Trends Hear 20. https://doi.org/10.1177/2331216516657466

Prendergast, G., Guest, H., Munro, K.J., Kluk, K., Léger, A., Hall, D.A., Heinz, M.G., Plack, C.J., 2017. Effects of noise exposure on young adults with normal audiograms I: Electrophysiology. Hear Res 344, 68–81. https://doi.org/10.1016/j.heares.2016.10.028

Prendergast, G., Tu, W., Guest, H., Millman, R.E., Kluk, K., Couth, S., Munro, K.J., Plack, C.J., 2018. Supra-threshold auditory brainstem response amplitudes in humans: Test-retest reliability, electrode montage and noise exposure. Hear. Res. https://doi.org/10.1016/j.heares.2018.04.002

Purcell, D.W., John, S.M., Schneider, B.A., Picton, T.W., 2004. Human temporal auditory acuity as assessed by envelope following responses. J. Acoust. Soc. Am. 116, 3581–3593. https://doi.org/10.1121/1.1798354

R Core Team, 2017. R: A language and environment for statistical computing. R Foundation for Statistical Computing, Vienna, Austria.

Ren, T., 2002. Longitudinal pattern of basilar membrane vibration in the sensitive cochlea. Proc. Natl. Acad. Sci. U.S.A. 99, 17101–17106. https://doi.org/10.1073/pnas.262663699

Russell, V., 2016. Least-Squares Means: The R Package lsmeans. Journal of Statistical Software 69, 1–33. https://doi.org/DOI:10.18637/jss.v069.i01

Schaette, R., McAlpine, D., 2011. Tinnitus with a normal audiogram: physiological evidence for hidden hearing loss and computational model. J. Neurosci. 31, 13452–13457. https://doi.org/10.1523/JNEUROSCI.2156-11.2011

Schmiedt, R.A., Mills, J.H., Boettcher, F.A., 1996. Age-related loss of activity of auditory-nerve fibers. J. Neurophysiol. 76, 2799–2803. https://doi.org/10.1152/jn.1996.76.4.2799

Sergeyenko, Y., Lall, K., Liberman, M.C., Kujawa, S.G., 2013. Age-related cochlear synaptopathy: an early-onset contributor to auditory functional decline. J. Neurosci. 33, 13686–13694. https://doi.org/10.1523/JNEUROSCI.1783-13.2013

Shaheen, L.A., Valero, M.D., Liberman, M.C., 2015. Towards a Diagnosis of Cochlear Neuropathy with Envelope Following Responses. J. Assoc. Res. Otolaryngol. 16, 727–745. https://doi.org/10.1007/s10162-015-0539-3

Stamataki, S., Francis, H.W., Lehar, M., May, B.J., Ryugo, D.K., 2006. Synaptic alterations at inner hair cells precede spiral ganglion cell loss in aging C57BL/6J mice. Hear. Res. 221, 104–118. https://doi.org/10.1016/j.heares.2006.07.014

Tichko, P., Skoe, E., 2017. Frequency-dependent fine structure in the frequency-following response: The byproduct of multiple generators. Hear. Res. 348, 1–15. https://doi.org/10.1016/j.heares.2017.01.014

Trune, D.R., Mitchell, C., Phillips, D.S., 1988. The relative importance of head size, gender and age on the auditory brainstem response. Hear. Res. 32, 165–174. https://doi.org/10.1016/0378-5955(88)90088-3

Valderrama, J.T., Beach, E.F., Yeend, I., Sharma, M., Van Dun, B., Dillon, H., 2018. Effects of lifetime noise exposure on the middle-age human auditory brainstem response, tinnitus and speech-in-noise intelligibility. Hear. Res. 365, 36–48. https://doi.org/10.1016/j.heares.2018.06.003

Valero, M.D., Burton, J.A., Hauser, S.N., Hackett, T.A., Ramachandran, R., Liberman, M.C., 2017. Noise-induced cochlear synaptopathy in rhesus monkeys (Macaca mulatta). Hear. Res. https://doi.org/10.1016/j.heares.2017.07.003

Verhulst, S., Altoè, A., Vasilkov, V., 2018a. Computational modeling of the human auditory periphery: Auditory-nerve responses, evoked potentials and hearing loss. Hear. Res. 360, 55–75. https://doi.org/10.1016/j.heares.2017.12.018

Verhulst, S., Ernst, F., Garrett, M., Vasilkov, V., 2018b. Supra-threshold psychoacoustics and envelope-following response relations: normal-hearing, synaptopathy and cochlear gain loss. Acta Acustica united with Acustica 104, 800–803.

Verhulst, S., Jagadeesh, A., Mauermann, M., Ernst, F., 2016. Individual Differences in Auditory Brainstem Response Wave Characteristics: Relations to Different Aspects of Peripheral Hearing Loss. Trends Hear 20, 1–20. https://doi.org/10.1177/2331216516672186

Voytek, B., Kramer, M.A., Case, J., Lepage, K.Q., Tempesta, Z.R., Knight, R.T., Gazzaley, A., 2015. Age-Related Changes in 1/f Neural Electrophysiological Noise. J. Neurosci. 35, 13257–13265. https://doi.org/10.1523/JNEUROSCI.2332-14.2015

Wu, P.Z., Liberman, L.D., Bennett, K., de Gruttola, V., O’Malley, J.T., Liberman, M.C., 2018. Primary Neural Degeneration in the Human Cochlea: Evidence for Hidden Hearing Loss in the Aging Ear. Neuroscience. https://doi.org/10.1016/j.neuroscience.2018.07.053

Yeend, I., Beach, E.F., Sharma, M., Dillon, H., 2017. The effects of noise exposure and musical training on suprathreshold auditory processing and speech perception in noise. Hearing Research. https://doi.org/10.1016/j.heares.2017.07.006

Zhong, Z., Henry, K.S., Heinz, M.G., 2014. Sensorineural hearing loss amplifies neural coding of envelope information in the central auditory system of chinchillas. Hear. Res. 309, 55–62. https://doi.org/10.1016/j.heares.2013.11.006

Zhu, L., Bharadwaj, H., Xia, J., Shinn-Cunningham, B., 2013. A comparison of spectral magnitude and phase-locking value analyses of the frequency-following response to complex tones. J. Acoust. Soc. Am. 134, 384–395. https://doi.org/10.1121/1.4807498

